# Endogenous mutational mechanisms and metabolic context shape endometrial cancer

**DOI:** 10.64898/2026.04.03.716356

**Authors:** Jian Sang, Mengyan Zhang, Sergio Chavez, Yewon Kim, Thomas Veith, Weiyin Zhou, Wen Luo, Adriana Morales Miranda, Jens Luebeck, Guangyu Wang, Bin Zhu, Vineet Bafna, Stephen J Chanock, Tongwu Zhang

## Abstract

Uterine corpus endometrial carcinoma (UCEC) is a common gynecologic malignancy with rising mortality, yet its genome-wide mutational architecture remains incompletely understood. Here we analyze deep whole-genome sequencing data from 440 UCEC tumors from The Cancer Genome Atlas, integrated with transcriptomic and clinical data, to define subtype-specific mutational processes and genomic architectures. We uncover pronounced molecular subtype-specific differences in endogenous mutational mechanisms, retrotransposition activity, structural variation, and chromosomal instability. LINE-1 retrotransposition emerges as a key contributor to genome instability in UCEC, acting as a prominent source of structural variation in copy-number stable tumors and associating with chromothripsis and ecDNA-mediated oncogene amplification in copy-number high tumors. Mutational signature SBS28 contributes significantly to mutations in *POLE*-deficient tumors and is strongly correlated with SBS10a and SBS10b. Mismatch repair-deficient tumors exhibit a previously unrecognized doublet base substitution signature and a defining imbalance between ID2 and ID1 indel processes, reflecting pervasive DNA template-strand replication slippage and associated with increased tumor immunogenicity. Copy-number low tumors follow a distinct low-mutagenesis evolutionary trajectory characterized by reduced replication stress, low proliferative activity, genomic stability and enrichment of SBS18 associated with oxidative damage. Notably, we identify a consistent inverse association between body mass index and tumor mutational burden in UCEC, suggesting that host metabolic state may influence fundamental cellular processes governing mutation accumulation, promoting tumor development through non-mutagenic mechanisms rather than elevated genomic instability. Together, these findings establish a mutagenesis-centric framework for UCEC that links endogenous mutational mechanisms to tumor architecture and host metabolic context, uncovering previously unrecognized subtype-specific genomic features with important implications for refined risk stratification and therapeutic strategies.

## Introduction

Uterine corpus endometrial carcinoma (UCEC) is among the most prevalent gynecologic malignancies worldwide and represents a growing public health concern, particularly in high-income countries. In the United States, approximately 68,270 new cases and 14,450 deaths are projected in 2026^1^. Despite substantial advances in prevention, early detection, and treatment, UCEC remains one of the few major cancers for which mortality shows a continued upward trend, even in settings with advanced healthcare systems. This growing burden is compounded by pronounced disparities in incidence and outcomes across population groups, revealing pervasive inequities in UCEC care and prognosis^2^. Epidemiological evidence supports a multifactorial etiology driven by prolonged unopposed estrogen exposure, obesity and metabolic dysfunction, reproductive factors, inherited genetic susceptibility, and aging^3^. Together, these factors contribute to the extensive biological heterogeneity observed in UCEC and highlight critical gaps in our understanding of the mechanisms driving tumor evolution and therapeutic response.

The Cancer Genome Atlas (TCGA) provides a foundational molecular framework that stratifies UCEC into four major subtypes: POLE-ultramutated (POLE), microsatellite instability-high (MSI), copy-number low (CN-Low), and copy-number high (CN-High), which reflects distinct patterns of genomic instability^4^. POLE and MSI tumors are characterized by exceptionally high mutation burdens arising from defects in DNA proofreading or mismatch repair, respectively, whereas CN-High tumors display extensive chromosomal instability and frequent *TP53* mutations. In contrast, CN-Low tumors harbor comparatively stable genomes^4,5^. This classification has substantially informed biological understanding and clinical risk stratification. However, this framework was derived primarily from whole-exome sequencing (WES), which limits resolution of genome-wide mutational processes and fails to capture key genomic features—such as large-scale structural variation, complex genomic rearrangements, transposable element activity, and extrachromosomal DNA (ecDNA) that are increasingly recognized as drivers of tumor aggressiveness, treatment response, and clinical outcome. Moreover, emerging evidence points to substantial heterogeneity within CN-Low and CN-High tumors, suggesting that current classifications capture only part of the underlying genomic complexity^6,7^. Beyond subtype-defining alterations, UCEC genomes are shaped by diverse endogenous mutational processes that operate genome-wide, giving rise to heterogeneous mutational signatures and complex patterns of genomic instability^8,9^. How these processes interact to generate subtype-specific genomic architectures, and how they relate to host factors such as obesity, remains incompletely understood. Addressing these gaps requires a comprehensive genome-wide approach that integrates high-depth whole-genome sequencing (WGS) with subtype-aware analysis and multi-omics profiling to systematically interrogate mutational processes and their structural consequences.

Here, we leverage deep WGS of 440 UCEC tumors from the TCGA cohort, together with matched transcriptomic and clinical data, to construct a comprehensive, genome-wide view of UCEC mutagenesis. By systematically characterizing diverse classes of genome-wide somatic variation, we delineate subtype-specific mutational processes that extend beyond existing WES-based frameworks. We identify LINE-1-associated structural rearrangements as a unifying driver of genomic instability across UCEC, link chromothripsis and ecDNA-mediated oncogene amplification to CN-High tumors, uncover a previously unrecognized mismatch repair-associated doublet base substitution (DBS) signature and a defining ID2-to-ID1 indel bias in MSI tumors, and reveal a striking inverse association between body mass index and tumor mutational burden that characterizes the low-proliferative, copy-number-stable CN-Low subtype. Together, our findings establish a genome-wide, mutagenesis-centric framework for understanding UCEC heterogeneity, linking endogenous mutational mechanisms to tumor architecture, evolutionary dynamics, and potential host metabolic state. This work provides a refined biological basis for molecular stratification and highlights distinct mutational vulnerabilities with implications for precision oncology in endometrial cancer.

## Results

### Distinct genomic architectures define UCEC subtypes

We analyzed deep WGS data from 440 UCEC tumors from TCGA, integrating matched transcriptomic and clinical data to systematically characterize subtype-specific mutational processes and structural variation (**Supplementary Data 1**). Median sequencing depths were 87× for tumors and 37× for matched blood or normal samples (**Supplementary Fig. 1a**). Tumor purity was generally high, with a median purity of 0.77 across UCEC tumors (**Supplementary Fig. 1b**). Using WGS-derived genomic features, we refined UCEC classification into four molecular subtypes that were highly concordant with the original TCGA whole-exome sequencing (WES) framework^4^ (92% concordance), with discrepancies arising between the CN-Low and CN-High groups. Specifically, we identified 41 POLE-ultramutated, 136 MSI, 173 CN-High, and 90 CN-Low tumors, each exhibiting a strikingly distinct genomic architecture (**Fig. 1**). Notably, tumors with serous-like histology were predominantly enriched within the CN-High subtype (Fisher’s exact test, P=7.08e-37) and were significantly associated with older age at diagnosis (two-sided Wilcoxon rank-sum test, P=2.24e-07; **Supplementary Fig. 1c-d).** These subtypes differed markedly in tumor mutational burden (TMB), proportion of genome altered (PGA), structural variant (SV) burden, and transposable element (TE) insertions (**Fig. 1b**). POLE tumors showed the highest single-nucleotide variant (SNV) burden, MSI tumors were enriched for indel mutations (**Fig. 1c**), CN-High tumors exhibited extensive copy-number alterations accompanied by elevated SV and TE activity, and CN-Low tumors displayed relatively stable genomes with low overall genomic alteration burden. Whole-genome duplication (WGD) was significantly enriched in CN-High tumors (59%; Fisher’s exact test, P=3.83e-27), consistent with their highly rearranged genomic profiles and elevated *TP53* mutations (**Fig. 1d**). In addition, CN-High tumors exhibited increased mitochondrial DNA (mtDNA) copy number compared with other subtypes (**Fig. 1e**), suggesting altered replicative or metabolic states associated with large-scale genomic instability.

**Fig. 1:**
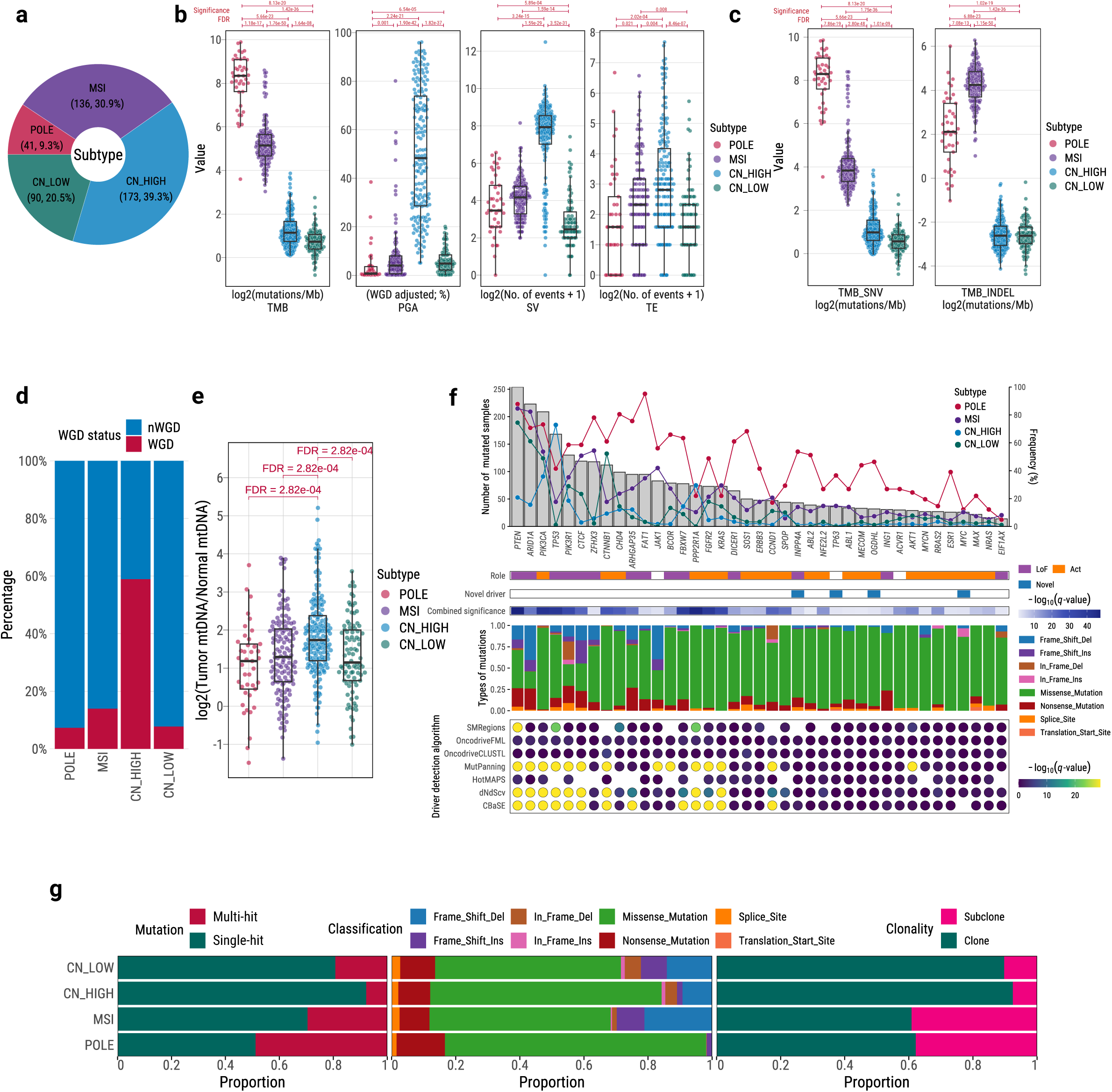
Whole-genome sequencing defines distinct genomic architectures across UCEC subtypes. **a**, Distribution of UCEC tumors classified into four molecular subtypes based on WGS-derived genomic features: POLE (n = 41), MSI (n = 136), CN-High (n = 173), and CN-Low (n = 90). **b**, Genome-wide alteration metrics across UCEC molecular subtypes, including tumor mutational burden (TMB; log_₂_ mutations per megabase), proportion of genome altered (PGA; WGD-adjusted), structural variant (SV) burden (log_₂_ number of events + 1), and transposable element (TE) insertion burden (log_₂_ number of events + 1). Each point represents one tumor. Boxes show median and interquartile range; whiskers denote 1.5× interquartile range. Two-sided Wilcoxon rank-sum tests with FDR correction were used. **c**, Genome-wide tumor mutational burden stratified by mutation class, showing separate contributions from single-nucleotide variants (SNVs) and small insertions and deletions (indels) across UCEC molecular subtypes (shown similarly to **b**). **d**, Fraction of WGD positive and WGD negative tumors within each molecular subtype. **e**, Relative mitochondrial DNA (mtDNA) copy number across UCEC subtypes, expressed as log_₂_(tumor/normal mtDNA ratio). Statistical significance was assessed using two-sided Wilcoxon rank-sum tests with FDR correction. **f**, Significantly mutated driver genes identified using an integrative driver discovery framework. The upper panel shows the number of tumors harboring mutations in each gene, with subtype-specific frequencies indicated. Middle annotations summarize inferred functional roles, novelty, and combined statistical significance (−log_₁₀_ q value). The lower panel shows mutation class composition and support across multiple driver detection methods. **g**, Subtype-specific properties of driver gene alterations, including the proportion of genes affected by single versus multiple-hit mutations, distribution of mutation classifications, and fraction of clonal versus subclonal events.

Using a systematic driver discovery framework^10^, we identified 39 significantly mutated driver genes in UCEC, exhibiting strong subtype-specific patterns of oncogenic alteration (**Fig. 1f**), most consistent with previous UCEC studies. These driver genes displayed markedly different mutation frequencies across subtypes, and a few candidate driver genes were first reported in UCEC, including *MYC*, *TP63*, *OGDHL*, and *INPP4A*. Notably, POLE tumors were uniquely enriched for multi-hit mutations affecting the same driver gene, reflecting their ultramutated phenotype (**Fig. 1g**). In contrast, frameshift indels within driver genes were significantly depleted in POLE tumors compared with MSI tumors (Fisher’s exact test, P=0.007). This observation is concordant with the substantially lower genome-wide indel burden in POLE tumors relative to MSI tumors (**Fig. 1c**). Clonal architecture analysis further revealed pronounced differences between subtypes. POLE and MSI tumors harbored a high proportion of subclonal mutations, indicating increased intratumoral heterogeneity and prolonged or ongoing mutagenesis, whereas CN-High tumors were characterized by fewer subclonal SNVs but extensive clonal copy-number alterations (**Supplementary Fig. 2**). Together, these results demonstrate that UCEC subtypes are defined by distinct genomic architectures in mutational processes, genomic instability, and tumor evolutionary dynamics.

### LINE-1-associated chromothripsis and ecDNA formation in CN-High tumors

To better characterize genome-wide rearrangements in UCEC, we analyzed the patterns of transposable element (TE) insertions and structural variants (SVs) across molecular subtypes. The vast majority (98%) of TE insertions were attributable to LINE-1 retrotransposition events (**Supplementary Data 2**), consistent with previous pan-cancer analyses^11^. Remarkably, the majority of somatic LINE-1 transductions with an identified germline source originated from a single germline LINE-1 element located at chr22q12.1 (48.8%), with a secondary contribution from chrXp22.2 (11.4%) (**Fig. 2a**). This pattern was consistent across subtypes (**Supplementary Fig. 3**). Notably, the germline LINE-1 element at chr22q12.1 has also been reported as the most frequently activated source element across cancer types^11^. These findings indicate that a limited number of highly active germline LINE-1 elements drive retrotransposition in UCEC. Interestingly, recurrent SV breakpoint analysis revealed a rearrangement hotspot dominated by translocation at chr22q12.1 in POLE, MSI, and CN-Low tumors—subtypes characterized by low copy-number alteration burden (**Fig. 2b**). This SV hotspot corresponds to the dominant germline LINE-1 source element and reflects LINE-1 insertion events at different target sites. In contrast, although CN-High tumors exhibited a substantially higher overall SV burden, their rearrangements were distributed more diffusely across the genome and did not form focal hotspots. Together, these observations suggest that LINE-1 activity is a major contributor to SV in low–copy-number UCEC subtypes, whereas CN-High tumors, which are enriched for *TP53* mutations (72.8%), exhibit widespread genomic instability in which additional mechanisms likely contribute to the diffuse structural variation landscape.

**Fig. 2:**
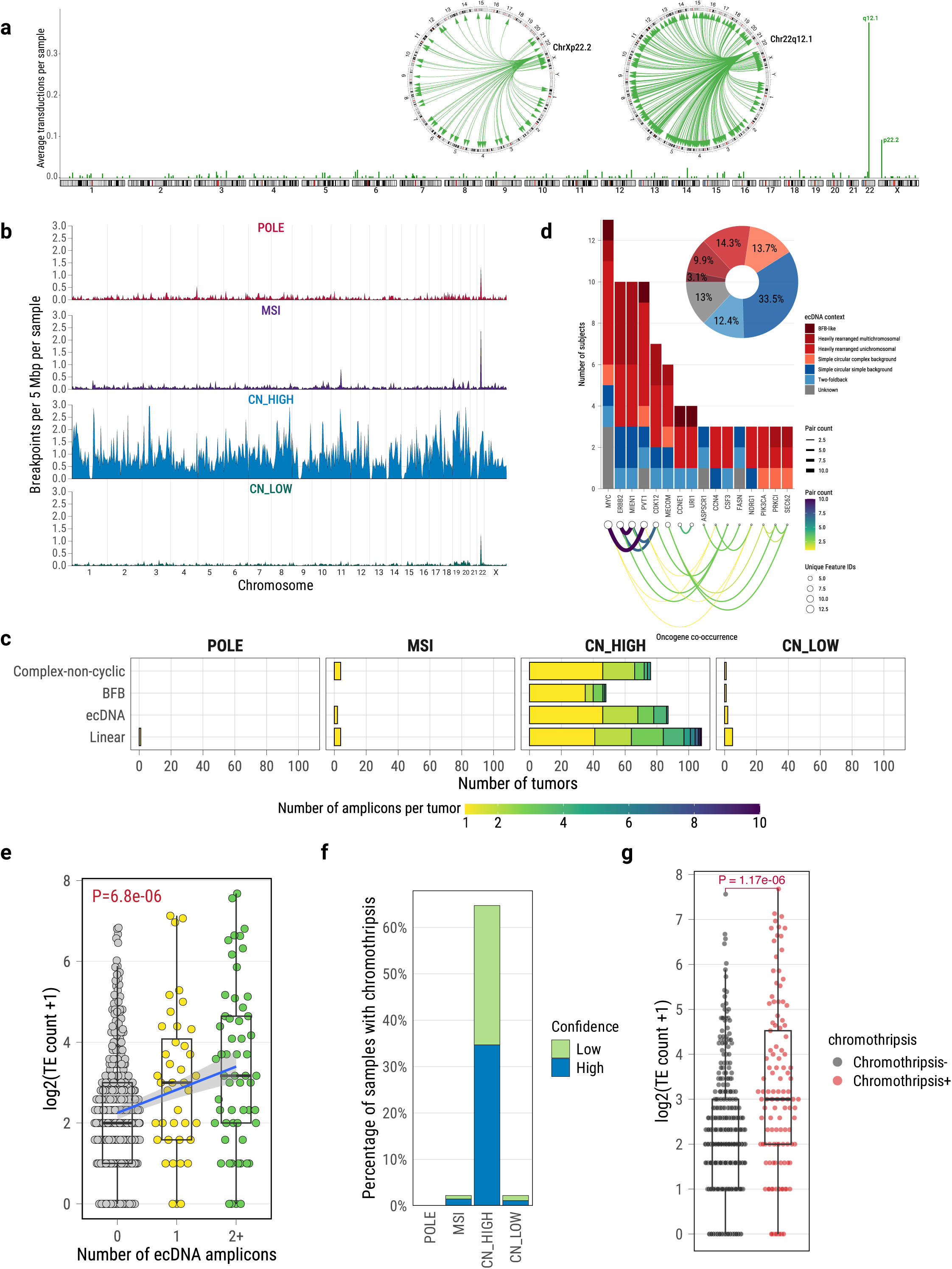
LINE-1 activity is associated with structural variation, chromothripsis, and ecDNA formation in UCEC. **a**, Genome-wide distribution of somatic LINE-1-mediated transductions across autosomes and sex chromosomes, shown as the average number of transduction events per sample in fixed genomic bins. Circos plots depict transduction events originating from recurrent germline LINE-1 source elements, including chr22q12.1 and chrXp22.2, with arcs indicating source-to-insertion relationships. **b**, Genome-wide profiles of structural variant (SV) breakpoints across UCEC molecular subtypes. SV breakpoint density is shown as the number of breakpoints per 5 Mb window, averaged across tumors within each subtype (POLE, MSI, CN-High, and CN-Low), and plotted along chromosomal coordinates. **c**, Summary of focal amplification architectures detected across molecular subtypes. Bars indicate the number of tumors harboring focal amplification events classified as linear, ecDNA, BFB, or complex non-cyclic structures, stratified by subtype. **d**, Characterization of oncogene-containing ecDNA amplicons. Stacked bar plots show the number of subjects harboring ecDNA-associated oncogenes, stratified by genomic context of ecDNA formation. Curved links illustrate pairwise co-occurrence of oncogenes detected on the same ecDNA amplicon, with line width and color corresponding to the number of shared events and node size indicating the number of subjects. **e**, Relationship between ecDNA burden and transposable element insertion counts. Tumors are grouped by the number of detected ecDNA amplicons (0, 1, or ≥2), and TE insertion burden is shown as log_₂_(TE count + 1). Boxes indicate median and interquartile range; whiskers represent 1.5× interquartile range. Statistical significance was assessed using a linear regression. **f**, Frequency of chromothripsis events across molecular subtypes. Stacked bars indicate the percentage of tumors with chromothripsis, stratified by confidence level (high or low) based on established detection criteria. **g**, Comparison of transposable element (TE) insertion burden between tumors with and without chromothripsis. TE burden is shown as log_₂_(TE count + 1) per tumor. Each dot represents an individual tumor. Boxes indicate median and interquartile range; whiskers represent 1.5× interquartile range. Statistical significance was assessed using a two-sided Wilcoxon rank-sum test.

To further investigate the relationship between LINE-1 activity and copy-number instability in CN-High tumors, we performed focal amplification and complex rearrangement analyses, including chromothripsis detection. Focal amplifications were observed predominantly in CN-High tumors (**Fig. 2c**), occurring in 79.8% of tumors, compared with 2.4% in POLE, 7.4% in MSI, and 8.9% in CN-Low tumors (**Supplementary Data 3**). All types of focal amplification mechanisms were significantly associated with high copy-number clusters or WGD in CN-High tumors (**Supplementary Fig. 4**). Notably, ecDNA was detected in 50.3% of CN-High tumors. Among ecDNA-positive tumors, 70.3% harbored at least one oncogene, with an average of 1.8 ecDNA amplicons per tumor. Classification of amplification architectures further revealed that ecDNA frequently arose within heavily rearranged chromosomal contexts rather than simple circular structures (**Fig. 2d**), and ecDNA-positive tumors showed markedly increased expression of amplified oncogenes (**Supplementary Fig. 5**). Tumors harboring ecDNA showed increased expression of cell cycle and proliferation-associated hallmark pathways, including *E2F targets*, *MYC targets*, and *G2M checkpoint* gene sets (**Supplementary Fig. 6**). Reconstruction of ecDNA amplicons revealed that *MYC-PVT1* and *ERBB2-MIEN1* were the most recurrent oncogene-containing ecDNA structures in CN-High tumors, occurring in 7.5% and 5.8% of tumors, respectively. Consistent with previous studies, *PVT1* represents one of the most frequently rearranged loci on ecDNA across cancer types, and ecDNA-associated *PVT1* fusions have been shown to stabilize and amplify oncogenic mRNAs^12,13^. Similarly, *ERBB2* (HER2) and *MIEN1*, co-localized on chromosome 17q12-21, were frequently co-amplified on ecDNA; *MIEN1* is a known oncogene that promotes cancer cell migration and invasion, and its co-amplification with *ERBB2* has been linked to aggressive tumor behavior^14–16^. Compared with ecDNA-negative tumors, ecDNA-positive tumors exhibited significantly worse overall survival (Cox proportional hazards model, P=2.15e-05) and progression-free interval (P=9.23e-07) (**Supplementary Fig. 7a-b**). However, within the CN-High subtype, no significant survival difference was observed between ecDNA-positive and ecDNA-negative tumors (**Supplementary Fig. 7c-d**). Among CN-High tumors, those harboring *MYC*-associated ecDNA showed a trend toward poorer overall survival and progression-free interval compared with other CN-High tumors (**Supplementary Fig. 7e-f**).

Importantly, within the CN-High subtype, tumors harboring a higher number of ecDNA amplicons exhibited a significantly increased LINE-1 insertion burden (**Fig. 2e**), consistent with previous reports linking LINE-1 activity to ecDNA formation^17–19^. Furthermore, 43.7% of ecDNA-positive tumors contained LINE-1 insertions on the same chromosome as the ecDNA, and 13.8% harbored LINE-1 insertions within 1 Mb of ecDNA breakpoints. In a small number of tumors (N = 12), LINE-1 insertions were detected within ecDNA structures themselves (**Extended Data Fig. 1a-b**). Together, these observations are consistent with a potential role for LINE-1-associated DNA breaks in facilitating amplicon formation, including ecDNA.

Chromothripsis, a catastrophic form of genomic rearrangement characterized by tens to hundreds of DNA double-strand breaks affecting one or multiple chromosomes, was observed almost exclusively in CN-High tumors, with a frequency of 64.7% (**Fig. 2f**; **Supplementary Data 4**). EcDNA-positive tumors were strongly associated with chromothripsis (Fisher’s exact test, P = 1.35e-08; **Supplementary Fig. 8**) within the CN-High subtype, suggesting a shared origin rooted in catastrophic chromosomal rearrangements. Chromothripsis-positive tumors also showed worse survival compared with other tumors (**Supplementary Fig. 9)**. Notably, LINE-1 insertion burden was significantly higher in tumors with chromothripsis than in those without (two-sided Wilcoxon rank-sum test; P=1.17e-06 across all tumors and P=0.05 within the CN-High subtype; **Fig. 2g**; **Supplementary Fig. 10**), further supporting an association between retrotransposition activity and catastrophic genomic rearrangements.

Collectively, these results indicate that LINE-1 retrotransposition is a major contributor to structural variation in copy-number-low UCEC tumors and is also associated with genomic instability in CN-High tumors, including chromothripsis and ecDNA-mediated oncogene amplification.

### Distinct mutational signatures define UCEC subtypes

Comprehensive mutational signature analysis revealed pronounced subtype specificity across single-base substitutions (SBS) and DBS (**Fig. 3a-c**; **Supplementary Data 5**). As expected, all POLE-ultramutated tumors exhibited strong contributions from SBS10a and SBS10b, reflecting exonuclease-domain mutations in DNA polymerase ε and accounting for the exceptionally high mutational burden in this subtype^20^. In 61% of POLE tumors, SBS10a and SBS10b together contributed more than 50% of all mutations. POLE tumors also showed enrichment of DNA mismatch repair deficiency related signatures, including SBS14, SBS15, SBS21, SBS26, and SBS44, detected collectively in 73% of POLE tumors with >5% mutation contribution. Consistent with the previous report^21^, SBS28 was strongly associated with SBS10a and SBS10b (Pearson correlation R=0.95, P=1.48e-20; **Extended Data Fig. 2a**), observed almost exclusively in POLE tumors (80.5%), and contributed substantially to the overall mutational burden (13.5%) in these tumors. Moreover, SBS28 exposure showed a negative association with SBS30, SBS32, and MSI-related signatures (including SBS15 and SBS44) (**Extended Data Fig. 3**), suggesting partially mutually exclusive mutational processes within POLE tumors. Together, these patterns suggest that SBS28 likely reflects a mutational process closely related to *POLE* exonuclease deficiency, sharing a similar underlying etiology with SBS10a and SBS10b.

**Fig. 3:**
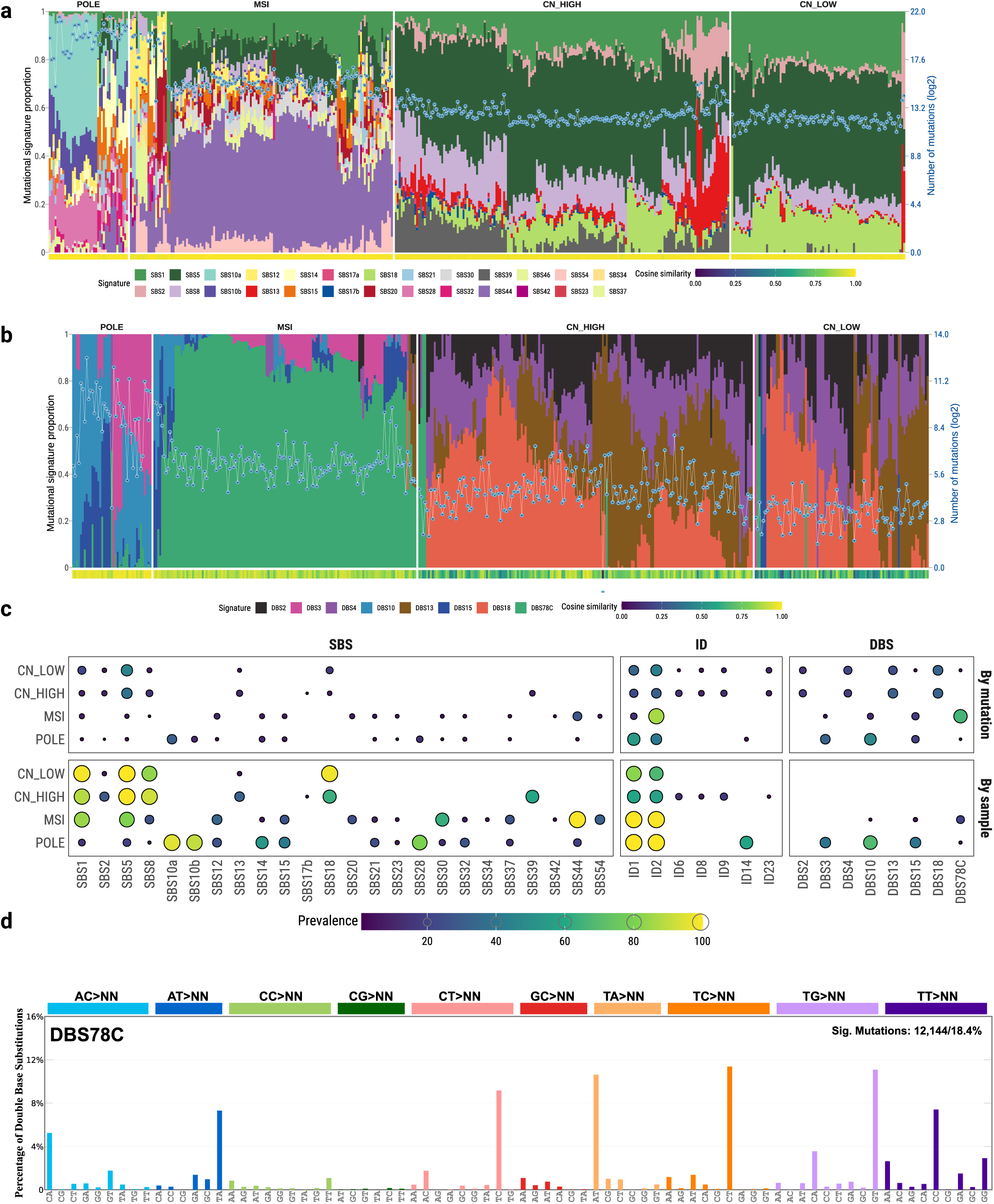
Subtype-specific mutational signature landscapes in UCEC. **a**, SBS mutational signature decomposition across UCEC tumors, ordered by molecular subtype. Stacked bars show the relative contribution of each SBS signature per tumor (left axis), with total SBS mutation counts shown as a line plot (right axis). Cosine similarities between the original and reconstructed mutational profiles are shown below. **b**, DBS mutational signature decomposition across UCEC tumors, displayed similarly to **a**, with stacked bars indicating relative signature contributions and total DBS mutation counts shown as a line plot. Cosine similarity values are shown below. **c**, Prevalence of SBS, ID, and DBS mutational signatures across UCEC molecular subtypes. Bubble size and color indicate the proportion of mutations (top) or the proportion of tumors (bottom) attributed to each signature within each subtype. A mutational signature was considered present in a tumor if at least 50 mutations were assigned to the signature and it contributed more than 5% of total mutations. **d**, *De novo* mutational signature profile of DBS78C identified in mismatch repair-deficient tumors, shown as relative contributions across doublet substitution contexts.

In the MSI subtype, SBS44 was the dominant mutational signature, present in nearly all tumors and accounting for 26.4% of mutations. Additional MSI-associated signatures, including SBS14, SBS15, SBS20, and SBS21, were also detected but contributed relatively fewer mutations (<8%). The reactive oxygen species-associated signature SBS18 was detected exclusively in CN-High and CN-Low tumors and was significantly enriched in CN-Low tumors compared with CN-High tumors (Fisher’s exact test; OR =25.0; P=1.43e-11), suggesting a distinct oxidative damage-related mutational process in copy-number-stable tumors.

APOBEC-mediated mutagenesis, represented by SBS2 and SBS13, was largely restricted to CN-High tumors, where it was detected in 36.4% of these tumors and contributed more than 20.4% of mutations, consistent with replication stress-associated cytidine deamination in this subtype. Within CN-High tumors, APOBEC signatures were significantly enriched in tumors harboring focal amplifications of any architecture (Fisher’s exact test; overall focal amplification OR=7.88; P=4.74e-07) and were also strongly associated with chromothripsis (OR=8.2; P=7.26e-10), suggesting that APOBEC activity preferentially arises in highly unstable genomic contexts and may contribute to the mutational diversification of genomically unstable CN-High tumors. Notably, SBS39, a mutational signature of unknown etiology, was almost exclusively observed in CN-High tumors with a prevalence of 59.5%. The strong enrichment of SBS39 in this subtype suggests a potential association with copy-number-driven genomic instability.

Given the high overall mutation burden in UCEC, we next examined DBS signatures. In POLE tumors, DBS10, DBS3, and DBS15 were the dominant DBS signatures, together accounting for 48.1%, 32.1% and 18.3% of DBS mutations. DBS10 was detected in nearly all POLE tumors. DBS3, previously associated with polymerase ε exonuclease-domain mutations, was identified in approximately 39% of POLE tumors and showed significant correlation with SBS28 signatures (Pearson correlation R=0.96, P=1.60e-15; **Extended Data Fig. 2b**), further supporting a shared mutational etiology between SBS28 and POLE exonuclease deficiency. The etiology of DBS15 remains unknown; however, its strong enrichment in POLE tumors suggests a potential association with polymerase ε exonuclease-domain mutations.

Strikingly, we identified a novel doublet base substitution signature, DBS78C, present in nearly all MSI tumors, where it accounted for 65.8% of DBS mutations, and in 51% of POLE tumors (based on DBS78C > 0; **Fig. 3d; Supplementary Data 6**). Mutations attributed to DBS78C were strongly correlated with estimated MSI scores and with the mutation burden of the major MSI-associated SBS signature SBS44 (Pearson correlation: R=0.72, P=2.44e-27; and R=0.61, P=1.65e-15, respectively; **Extended Data Fig. 4**). These results suggested that DBS78C represents a mutational process active in mismatch repair-deficient UCEC. Interestingly, DBS78C is characterized by a high frequency of A/T-containing reversed doublet substitutions (XY→YX). These events are consistent with replication slippage or template realignment in the absence of functional mismatch repair, potentially compounded by error-prone processing of tandem DNA lesions. To our knowledge, this pattern of reversed doublet substitutions has not been reported in existing mutational signature catalogs, suggesting that DBS78C represents a previously unrecognized MMR-associated doublet mutational process.

In contrast, CN-Low and CN-High tumors exhibited substantially lower DBS mutation burdens than MSI and POLE tumors, with an average of 11.4, 35.6, 119, and 1,042 DBS mutations per tumor in CN-Low, CN-High, MSI, and POLE tumors, respectively. The overall DBS mutational landscapes of CN-Low and CN-High tumors were broadly similar; however, the low number of DBS events in these subtypes limited the statistical power to robustly identify subtype-specific DBS signatures. In addition, distinct patterns of copy-number and structural variant signatures were observed across the four UCEC subtypes, including the identification of novel SV signatures enriched in CN-High tumors (e.g., SV32A; **Extended Data Fig. 5; Supplementary Data 7-8**).

### Elevated ID2-to-ID1 ratio characterizes MSI tumors

Analysis of indel mutational signatures revealed pronounced subtype-specific patterns across UCEC (**Fig. 4a**). Replication slippage-associated signatures ID1 and ID2 dominated the indel landscape in all subtypes^22^, together accounting for 95.8%, 100%, 55.0%, and 75.9% of indels in POLE, MSI, CN-High, and CN-Low tumors, respectively. ID14, a signature of unknown etiology, was detected exclusively in 58.5% of POLE tumors. CN-High and CN-Low tumors exhibited broadly similar indel signature profiles. However, CN-High tumors showed significant enrichment of ID6 (associated with defective homologous recombination), ID8 (associated with non-homologous end joining), and ID9 (unknown etiology), accompanied by relative depletion of ID1 and ID2 signatures compared with CN-Low tumors (**Supplementary Fig. 11**). This pattern is consistent with increased genomic instability-associated indel mutagenesis in CN-High tumors.

**Fig. 4:**
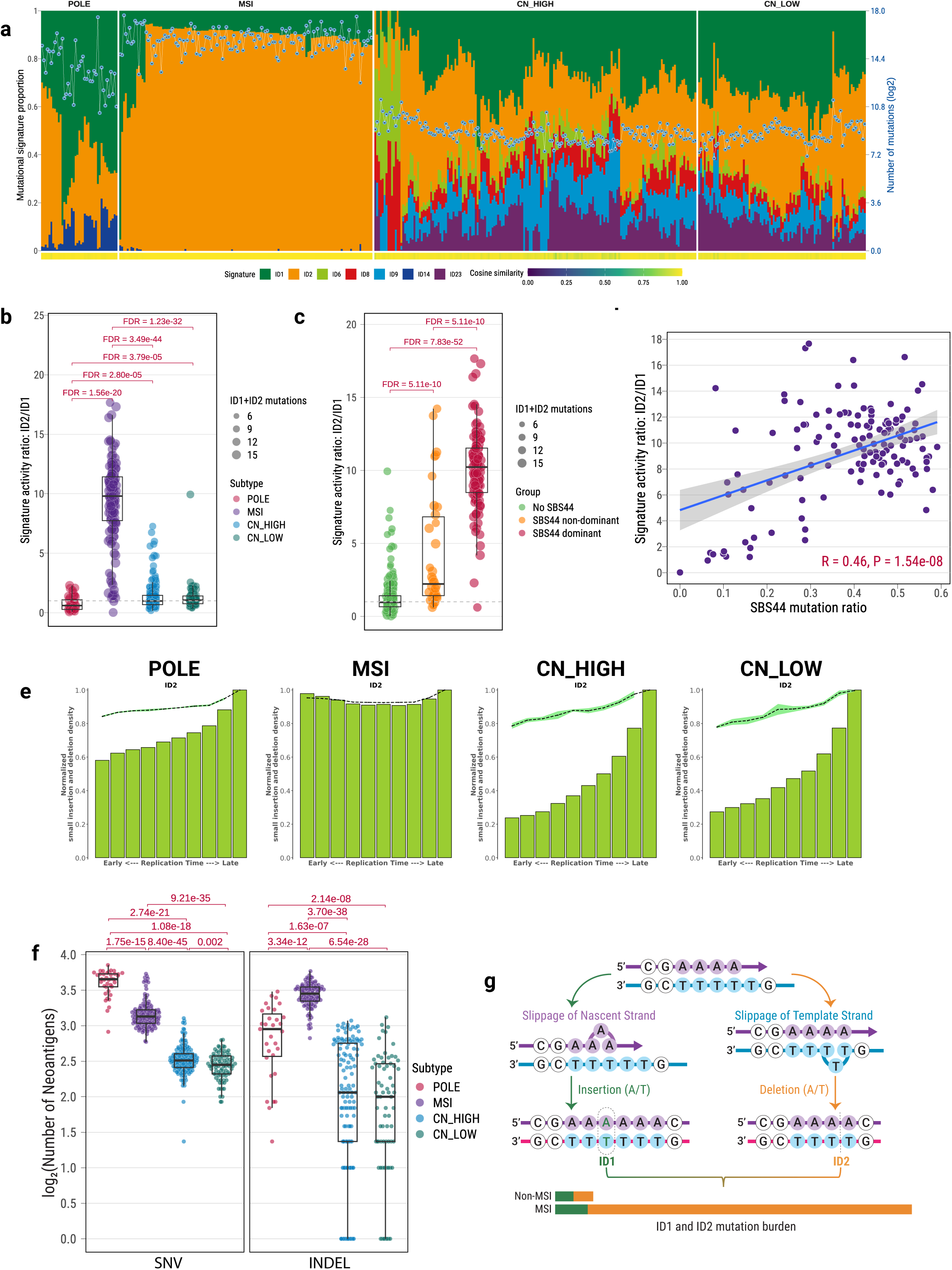
Indel mutational signatures distinguish UCEC molecular subtypes. **a**, ID mutational signature decomposition across UCEC tumors, ordered by molecular subtype. Stacked bars show the relative contribution of each ID signature per tumor (left axis), with total ID mutation counts shown as a line plot (right axis). Cosine similarities between the original and reconstructed ID profiles are shown below. **b**, Comparison of ID2-to-ID1 signature activity ratios across UCEC molecular subtypes. Each point represents an individual tumor. Boxplots indicate the median and interquartile range. Statistical significance was assessed using two-sided Wilcoxon rank-sum tests with FDR correction. **c**, ID2-to-ID1 ratios stratified by dominance of the MSI-associated single-base substitution signature SBS44. Tumors were grouped as SBS44-dominant (>50% contribution), SBS44 non-dominant, or SBS44-absent. Statistical significance was assessed using two-sided Wilcoxon rank-sum tests with FDR correction. **d**, Correlation between ID2-to-ID1 ratio and SBS44 mutation burden across UCEC tumors. Pearson correlation coefficient (R) and two-sided P value are shown; the shaded region denotes the 95% confidence interval of the fitted linear regression. **e**, Replication timing distribution of ID2 mutations across UCEC molecular subtypes. Bars indicate normalized densities of indel mutations across replication timing bins from early to late replicating regions; dashed lines represent smoothed trends. **f**, Comparison of indel-derived neoantigen burden across UCEC molecular subtypes. Each point represents an individual tumor. Boxplots summarize distributions, with statistical significance assessed using two-sided Wilcoxon rank-sum tests and Benjamini-Hochberg correction; adjusted P values are shown. **g**, Schematic illustration of replication slippage mechanisms underlying ID1 and ID2 mutations. ID1 mutations arise from slippage of the nascent strand, whereas ID2 mutations result from slippage of the template strand. The relative contributions of ID1 and ID2 in MSI and non-MSI tumors are summarized.

Although overall indel burden was higher in both POLE and MSI tumors than in CN-High and CN-Low tumors, the relative contributions of ID1 and ID2 differed markedly between POLE and MSI subtypes (**Supplementary Fig. 12**). MSI tumors exhibited a highly distinctive indel landscape dominated almost exclusively by ID1 and ID2, with near-complete absence of other indel processes. While the ID1 mutation burden was comparable between POLE and MSI tumors, ID2 mutations were substantially more abundant in MSI tumors despite the much higher SNV burden in POLE tumors (**Extended Data Fig. 6**). Consequently, the ID2-to-ID1 ratio was markedly elevated in MSI tumors compared with other subtypes (**Fig. 4b**), showing an approximately ten-fold increase relative to POLE, CN-High, and CN-Low tumors. This striking shift in ID2/ID1 balance suggests a fundamental alteration in replication slippage dynamics associated with mismatch repair (MMR) deficiency.

Stratifying tumors by the dominant MSI-associated single-base substitution signature SBS44 further reinforced this association. MSI tumors with SBS44 contributing more than 50% of total mutations exhibited significantly higher ID2-to-ID1 ratios than tumors without dominant SBS44 activity (**Fig. 4c**). Moreover, the ID2-to-ID1 ratio showed a strong positive correlation with SBS44 mutation burden (Pearson correlation R=0.46, P=1.54e-08; **Fig. 4d**), indicating that increased ID2 bias tightly tracks the degree of MMR deficiency. Together, these results identify the ID2-to-ID1 ratio as a robust marker of MSI status.

Analysis of replication timing further revealed a unique topographical property of ID2 mutations in MSI tumors (**Fig. 4e**). In POLE, CN-High, and CN-Low tumors, ID2 mutations were significantly enriched in late-replicating genomic regions, consistent with patterns observed across many cancer types. In contrast, ID2 mutations in MSI tumors showed no enrichment across replication timing, indicating that ID2 mutagenesis in MSI occurs throughout the replication program. By comparison, ID1 mutations were consistently enriched in late-replicating regions across all UCEC subtypes (**Supplementary Fig. 13**). These findings suggest that, in the absence of functional mismatch repair, template-strand replication slippage (ID2) accumulates genome-wide and continuously during DNA replication.

To validate the specificity of this ID2 bias, we reanalyzed published pan-cancer whole-genome sequencing data from PCAWG^22^. While most tumor types exhibited relatively balanced ID1 and ID2 activity, a pronounced bias toward ID2 was observed in MSI-prone cancer types, including colorectal, stomach, and uterine cancers **(Extended Data Fig. 7a-b)**. Consistently, reanalysis of indel mutational signatures derived from CRISPR-Cas9-based knockouts of DNA repair genes^23^, identified a single ID signature with high similarity to ID2 (cosine similarity = 0.97; **Extended Data Fig. 7c**). Moreover, across both MMR-deficient backgrounds (*MSH6* and *EXO1*), the 1-bp T deletions (ID2) remained significantly higher than the 1-bp T insertions (ID1) (**Extended Data Fig. 7d)**, further supporting a strong bias toward ID2 associated with DNA mismatch repair deficiency. Functionally, the elevated ID2 burden in MSI tumors was associated with a significantly increased number of indel-derived neoantigens compared with other UCEC subtypes (**Fig. 4f**), suggesting enhanced immunogenic potential. Mechanistically, ID2 mutations arise from slippage of the template strand during DNA replication, whereas ID1 mutations reflect slippage of the nascent strand (**Fig. 4g**). Our results indicate that MMR deficiency selectively permits the accumulation of template-strand slippage events across the genome, leading to a strong ID2 bias.

Collectively, these findings demonstrate that an elevated ID2-to-ID1 ratio is a defining feature of MSI tumors and reflects a pervasive replication slippage mechanism driven by mismatch repair deficiency, with potential implications for tumor immunogenicity and clinical stratification.

### Higher body mass index is associated with lower TMB

To further understand the biological characteristics of CN-Low tumors, which exhibit substantially lower mutagenesis compared with POLE, MSI, and CN-High subtypes, we examined replication stress, proliferative activity, and clinical correlates across UCEC subtypes. CN-Low tumors displayed the lowest replication stress scores among all subtypes (**Fig. 5a**), accompanied by significantly reduced expression of proliferation-related gene signatures (**Fig. 5b**). Consistently, canonical proliferation markers, including *MKI67* and *TOP2A*, were expressed at significantly lower levels in CN-Low tumors compared with other subtypes (**Supplementary Figure 14a-b**). In addition, *PD-L1* expression was lowest in CN-Low tumors relative to other molecular subtypes (**Supplementary Figure 14c**), and CN-Low tumors exhibited significantly longer telomere length compared with CN-High tumors (**Supplementary Figure 15**). In contrast, expression of *XIST*, a marker of X-chromosome inactivation, was highest in CN-Low tumors (**Fig. 5c**), suggesting enhanced chromatin silencing and reduced replicative activity. Together, these features indicate that CN-Low tumors are characterized by a low-proliferative, low-replication stress cellular state. A previous study reported the presence of the *Armadillidium vulgare* iridescent virus (IIV31) in more than 25% of UCEC tumors^24^. Although the underlying mechanism remains unclear, we observed a strong enrichment of IIV31 in tumors from the CN-Low subtype (Fisher’s exact test; OR=3.0, P=1.60e-05; **Supplementary Figure 16**).

**Fig. 5:**
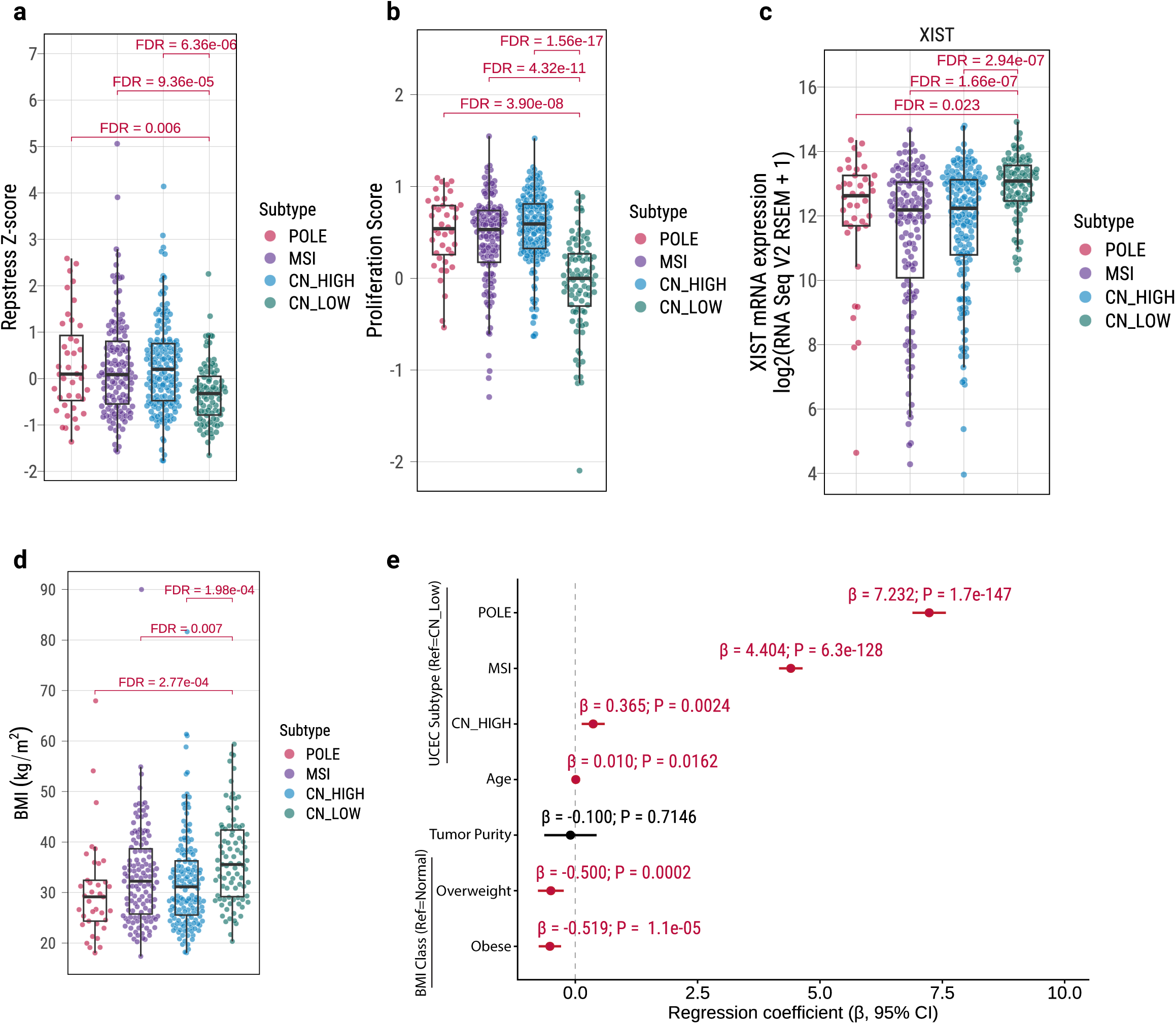
Transcriptomic features of CN-Low tumors and their association with body mass index. **a**, Replication stress Z-scores across UCEC molecular subtypes. Each point represents an individual tumor. Boxplots indicate the median and interquartile range. Pairwise statistical comparisons were performed using two-sided Wilcoxon rank-sum tests with FDR correction. **b**, Proliferation scores across UCEC molecular subtypes, derived from proliferation-related gene expression signatures. Each point represents one tumor. Boxplots summarize the distribution within each subtype. FDR-adjusted P values are shown. **c**, Expression of *XIST* across UCEC molecular subtypes. Each point represents one tumor. Boxplots indicate median and interquartile range. FDR-adjusted P values for pairwise comparisons are shown. **d**, Distribution of body mass index (BMI) across UCEC molecular subtypes. Each point represents an individual patient. Boxplots summarize the distribution within each subtype. FDR-adjusted P values for pairwise comparisons are indicated. **e-f**, Multivariable regression analysis assessing associations with tumor mutational burden (TMB) in TCGA-UCEC (**e**; *N*=409; TMB=log2(mutations/Mb)). Variables include UCEC molecular subtype (reference = CN-Low), age at diagnosis, tumor purity, and BMI class (reference = normal). Points represent regression coefficients (β), and horizontal lines indicate 95% confidence intervals. Two-sided P values are shown.

Interestingly, patients with CN-Low tumors exhibited significantly higher BMI compared with patients in other molecular subtypes, with 72.4% of CN-Low tumors arising in obese patients (BMI ≥ 30kg/m²; **Fig. 5d**; **Supplementary Fig. 17**). Integration of clinical and genomic variables revealed a significant inverse association between BMI and tumor mutational burden (TMB) (overall P=7.44e-05; multivariable regression adjusting for age at diagnosis, tumor purity, and UCEC molecular subtype; **Fig. 5e**). Notably, this negative association between BMI and TMB was consistently observed across all molecular subtypes (**Extended Data Figure 8**). We next examined associations between BMI and mutational processes adjusting for age at diagnosis, tumor purity, and molecular subtype. After correction for multiple testing, no mutational signatures were significantly positively associated with higher BMI. However, a borderline association was observed between SBS8 (of unknown etiology) and higher BMI (FDR=0.06; **Supplementary Fig. 18a**). In addition, no significant associations were detected between BMI and driver gene mutation status (**Supplementary Fig. 18b**).

Collectively, these findings indicate that higher BMI is associated with reduced nucleotide-level mutagenesis in UCEC, independent of molecular subtype. In particular, CN-Low tumors are characterized by a distinctive biological state marked by low replication stress, reduced proliferative activity, enhanced X-chromosome inactivation, longer telomere length, and enrichment in obese patients.

## Discussion

In this study, we provide a comprehensive whole-genome view of UCEC that reframes tumor heterogeneity through the lens of genome-wide mutational processes. By integrating deep whole-genome sequencing with transcriptomic and clinical data across 440 tumors, we extend the TCGA molecular framework beyond coding regions and demonstrate how distinct endogenous mutational mechanisms shape subtype-specific genomic architectures, tumor evolution, and clinical phenotypes.

Our analyses highlight an important role for LINE-1 retrotransposition in shaping genome instability across UCEC. *TP53* is known to suppress LINE-1 activity^26–28^, and in low–copy-number UCEC subtypes, where *TP53* mutations are infrequent, LINE-1 retrotransposition appears constrained by intact genome surveillance. Consistent with this, these tumors exhibit limited structural variation and lack widespread copy-number alterations. The shared focal SV hotspot with the dominant germline LINE-1 source locus indicates that genomic rearrangements in these subtypes largely reflect LINE-1 insertion events. In contrast, CN-High tumors are enriched for *TP53* mutations and display elevated LINE-1 insertion burden together with pervasive genome-wide structural variation. In this context, LINE-1–associated DNA breaks coincide with extensive chromosomal instability, chromothripsis, and frequent focal amplifications, including extrachromosomal DNA (ecDNA), a genomic architecture increasingly recognized as a driver of oncogene amplification^29^. The association between LINE-1 insertion burden, chromothripsis, and ecDNA formation suggests that retrotransposition-associated DNA damage may contribute to catastrophic genomic rearrangements from which ecDNA may emerge under strong oncogenic selection.

At the nucleotide level, our study refines the mutational landscapes of POLE and MSI tumors. In POLE-ultramutated tumors, mutagenesis is overwhelmingly dominated by single-nucleotide variants, with minimal contribution from indels, consistent with error-prone base substitution driven by polymerase ε exonuclease deficiency. We confirm the dominance of SBS10a and SBS10b and demonstrate that SBS28 is tightly linked to *POLE* exonuclease deficiency, contributing substantially to overall mutational burden and exhibiting mutually exclusive relationships with mismatch repair-associated signatures. Notably, frameshift indels within cancer driver genes were significantly depleted in POLE tumors, indicating that POLE-driven mutagenesis preferentially generates nucleotide substitutions rather than replication slippage-associated indels. Together, these findings highlight a distinct mutational regime in POLE tumors characterized by pervasive SNV accumulation with limited indel involvement in tumor evolution.

In MSI tumors, we identify a previously unrecognized doublet base substitution signature, DBS78C, which is strongly correlated with MSI burden and characterized by a high frequency of A/T-containing reversed doublet substitutions. This pattern likely reflects mismatch repair-dependent failure to resolve tandem replication errors and expands the repertoire of mutational processes associated with MMR deficiency. Indel mutagenesis further distinguished MSI tumors through a striking imbalance between ID2 and ID1 signatures. MSI tumors exhibited an approximately ten-fold increase in the ID2-to-ID1 ratio, driven predominantly by enrichment of 1-bp T deletions relative to insertions, reflecting pervasive template-strand replication slippage in the absence of functional mismatch repair. This bias was genome-wide, independent of replication timing, and consistently recapitulated across MSI-prone cancer types and CRISPR-based MMR knockout models. Importantly, the ID2-to-ID1 ratio relies on simple indel classes that are readily detectable from whole-exome sequencing and even targeted sequencing panels, highlighting its potential practical utility. Unlike other indel-enriched genomic instability states, such as homologous recombination deficiency, which generate more diverse indel spectra, the pronounced ID2 bias provides a mechanistically specific and robust marker of mismatch repair deficiency. The elevated ID2 burden was also associated with increased indel-derived neoantigens, providing a mechanistic link between replication slippage, tumor immunogenicity, and the clinical responsiveness of MSI tumors to immune checkpoint blockade.

In contrast, CN-Low tumors emerged as a biologically distinct, low-mutagenesis state. These tumors exhibited low replication stress, reduced proliferative activity, enhanced X-chromosome inactivation, longer telomere length, and minimal structural disruption. Strikingly, CN-Low tumors were enriched among obese patients. Importantly, body mass index (BMI) showed a strong inverse association with tumor mutational burden, a relationship that was consistent across all UCEC molecular subtypes. Although obesity is a well-established risk factor for UCEC, endometrial tumors lack a distinct BMI-associated mutational signature. Together, these findings suggest that obesity does not drive tumorigenesis through direct mutagenesis, but rather through systemic metabolic and inflammatory processes. This supports a model in which obesity promotes UCEC development via hormonal and metabolic alterations^30^, such as hyperestrogenism, insulin resistance, and chronic inflammation, without necessitating increased genomic instability, and highlights a fundamentally different mode of cancer risk compared to canonical mutagenic exposures like smoking. Reduced replication stress and proliferative dynamics in tumors arising in obese patients may limit replication-associated DNA damage and mutation accumulation, resulting in comparatively stable genomes. Alternatively, obesity-associated conditions may preferentially select for tumor clones with lower genomic instability rather than directly increasing mutation rates. The consistency of the BMI-mutational burden relationship across molecular subtypes suggests that host metabolic state potentially influences fundamental processes governing mutation accumulation, thereby shaping distinct evolutionary trajectories in UCEC.

These findings suggest that obesity drives endometrial cancer through systemic metabolic and inflammatory processes rather than direct mutagenesis, highlighting a fundamentally different mode of cancer risk compared to canonical exposures like smoking.

In summary, this study establishes a genome-wide, process-centric framework for understanding UCEC heterogeneity by linking endogenous mutational mechanisms to subtype-specific genomic architectures and host metabolic context. Our findings identify LINE-1-associated structural instability and ecDNA formation as key features of copy-number-driven disease, define mechanistically grounded mutational markers of mismatch repair deficiency, and reveal an obesity-associated, low-mutagenesis evolutionary trajectory in CN-Low tumors. Together, these results refine molecular stratification in UCEC and underscore the clinical value of the whole-genome approach for identifying biologically meaningful biomarkers and therapeutic vulnerabilities beyond mutation burden alone.

## Methods

### Multi-omics data collection, processing, and quality control

WGS data were obtained from 482 treatment-naive primary UCEC tumors with matched normal tissue (n = 424) or blood samples (n = 16). Raw WGS data (BAM files) together with clinical annotations were obtained from the Genomic Data Commons (GDC) data portal (https://portal.gdc.cancer.gov/). RNA sequencing (RNA-seq) data were obtained as preprocessed gene expression matrices for 529 UCEC tumors from the cBioPortal PanCancer Atlas cohort (https://www.cbioportal.org/), of which 427 tumors had matched WGS data. Clinical annotations for overall survival (OS) and progression-free interval (PFI) were obtained from the TCGA Pan-Cancer Atlas clinical dataset^31^. A total of 439 TCGA-UCEC patients with matched WGS data were included in downstream survival analyses. Additional clinical variables, including body mass index (BMI), were retrieved for the TCGA-UCEC cohorts from the cBioPortal PanCancer Atlas (https://www.cbioportal.org/). Among the TCGA-UCEC patients with available BMI information, 409 had matched WGS data and were included in integrative analyses. Patients were categorized according to World Health Organization criteria as normal weight (BMI 18.5-24.9 kg/m²), overweight (BMI 25.0-29.9 kg/m²), or obese (BMI ≥ 30 kg/m²).

Rigorous quality control was applied to all WGS samples prior to downstream analyses. Sequencing quality metrics were computed using Picard (v2.23.3), and genome-wide sequencing depth was quantified using mosdepth (v0.3.3). Tumor-normal sample concordance was evaluated using Somalier (v0.2.6), and genetic ancestry and potential sample contamination were assessed using VerifyBamID (v2.0.1). Tumor purity was estimated using the NGSpurity pipeline^32,33^. Samples were excluded if they failed any of the following predefined criteria: (i) median sequencing depth <20x for normal samples or <30x for tumor samples; (ii) tumor-normal concordance <0.9 based on homozygous variant comparison; (iii) estimated contamination (VerifyBamID FREEMIX) >0.05; or (iv) tumor purity <0.2. In addition, all tumors were required to pass automated quality control metrics and manual inspection of copy-number profiles generated by the NGSpurity pipeline. In total, 42 tumor-normal pairs failed quality control and were excluded from all downstream analyses.

To ensure uniform downstream processing, all BAM files were converted to FASTQ format and realigned to the GRCh38 human reference genome using a previously established unified preprocessing workflow^34^. This pipeline preserves read group information, sequencing lane metadata, and unmapped reads. Aligned reads were subsequently processed with duplicate marking, local realignment around small insertions and deletions, and base quality score recalibration. Final analysis-ready files were generated in compressed CRAM format and used for all downstream analyses.

### Somatic mutation calling and driver gene identification

Somatic single-nucleotide variants (SNVs) and small insertions and deletions (indels) were identified from analysis-ready CRAM files using our previously established deep WGS analysis pipeline^32,34–37^. Briefly, four complementary somatic variant callers Strelka (v2.9.10), MuTect, MuTect2, and TNscope were applied to each tumor-normal pair. MuTect, MuTect2, and TNscope were executed using the Sentieon Genomics framework (v202503). Variant calls from individual algorithms were subsequently integrated using an ensemble strategy designed to prioritize high-confidence somatic events. Candidate somatic variants were required to satisfy multiple quality-filtering criteria, including minimum supporting read counts, variant allele fraction thresholds, and exclusion of potential germline contamination. For indel detection, only variants independently identified by at least three callers were retained. All retained indels were left-normalized to ensure consistent representation across samples. Functional annotation of somatic variants was performed using ANNOVAR (v2020-06-08).

Driver gene discovery was performed using the IntOGen pipeline (v2020.02.0123)^10^, which integrates multiple orthogonal statistical approaches to identify genes under positive selection based on mutational recurrence, clustering, and functional impact. Genes achieving significance at a combined q-value < 0.1 were considered cohort-level driver genes. The mode of action for each driver gene (oncogene or tumor suppressor gene) was assigned based on the relative enrichment of truncating versus nonsynonymous mutations estimated by *dNdScv*^38^, together with annotations from the Cancer Gene Census. Genes with discordant evidence were classified as ambiguous.

### Allele-specific somatic copy number analysis

Allele-specific somatic copy-number alterations (SCNAs) were inferred from tumor-normal whole-genome sequencing data using a joint modelling framework that integrates tumor purity, ploidy, and clonal architecture, which was implemented within the NGSpurity pipeline^32,33^. Initial SCNA profiles were generated using Battenberg (v2.2.9), together with DPClust (v2.2.8) and ccube (v1.0). SCNA solutions were subjected to manual quality assessment, and profiles that did not meet predefined quality criteria were iteratively refitted by adjusting purity and ploidy parameters, informed by local copy-number states and subsequent manual review. Recurrent focal copy-number alterations were identified using GISTIC2.0 based on the major clonal copy-number state of each genomic segment.

### Structural variant and transposable element insertion analysis

Structural variants (SVs) were detected using two complementary algorithms, Meerkat (v0.189)^39^ and Manta (v1.6.0)^40^, each applied with recommended filtering parameters. A unified SV call set was generated by merging high-confidence calls from both methods. SVs were annotated for predicted effects on protein-coding genes, and candidate gene fusion events were retained based on established criteria, including gene-gene involvement, head-tail orientation, and preservation of coding frame.

Somatic transposable element (TE) insertions were identified using TraFiC-mem (v1.1.0)^11^. Only TE insertion events passing default quality and filtering criteria were retained for downstream analyses. Identified LINE-1 (L1) retrotransposition events were further classified according to their inferred source elements as germline-derived or somatic-derived^35^.

### UCEC molecular subtype classification

Tumor samples were classified into four molecular subtypes following the established TCGA integrative classification framework^4^: POLE-ultramutated, microsatellite instability-high (MSI), copy-number high (CN-High) and copy-number low (CN-Low). Initial subtype annotations were obtained from previously published TCGA studies and were based on integrated analyses of somatic mutation profiles, microsatellite instability status, and genome-wide copy-number alteration patterns. We applied a consistent and refined classification strategy to the samples analyzed in this study, leveraging high-resolution whole-genome sequencing data. Briefly, tumors were classified as POLE-ultramutated if they harbored pathogenic exonuclease-domain mutations in POLE. In addition, mutational signature analysis was required to show enrichment of POLE-associated signatures, with the combined contribution of SBS10a and SBS10b exceeding 5% of total mutations, corresponding to a minimum of 58,023 SBS10a/b-attributed mutations in our dataset.

Tumors were classified as MSI if they exhibited high levels of microsatellite instability. Specifically, MSI tumors were required to show enrichment of MSI-associated mutational signatures (SBS44, SBS14, SBS15, and SBS21) contributing more than 5% of total mutations, corresponding to a minimum of 4,566 MSI-attributed mutations in our dataset. In addition, tumors were required to have an MSI score greater than 2% as estimated by MSIsensor-pro^41^ and not be classified as POLE-ultramutated.

The remaining tumors were stratified into CN-Low and CN-High subtypes based on the extent and pattern of somatic copy-number alterations (**Supplementary Fig. 2**). Tumors exhibiting few genome-wide copy-number alterations and clustering within copy-number clusters 2 and 4 were classified as CN-Low, whereas tumors displaying extensive copy-number alterations and clustering within copy-number clusters 1 and 3 were classified as CN-High.

### Extrachromosomal DNA analysis

ecDNA was identified using the AmpliconSuite pipeline (v1.5.0). Briefly, tumor BAM files were processed to generate copy-number profiles and detect seed intervals, followed by reconstruction of amplified genomic structures using AmpliconArchitect (v1.5.r0). Reconstructed amplicons were subsequently classified using AmpliconClassifier (v1.5.1) into ecDNA or alternative amplification architectures, including breakage-fusion-bridge (BFB) cycles, linear amplifications, and complex non-cyclic events. Candidate ecDNA structures were further subjected to manual visual inspection to confirm classification and exclude ambiguous cases.

### Chromothripsis inference

Chromothripsis was inferred from whole-genome sequencing data using ShatterSeek^42^. Allele-specific copy-number profiles and structural variant calls were provided as input. Chromothripsis-positive chromosomes were identified based on established multivariable decision criteria implemented in ShatterSeek, following the high-confidence definitions used by the PCAWG study. Briefly, chromosomes were classified as high-confidence chromothripsis events if they satisfied one of the following criteria: (i) at least six interleaved intrachromosomal SVs, seven or more contiguous copy-number segments oscillating between two states, failure of the fragment-joins test, and a significant chromosomal enrichment or exponential distribution of breakpoints test (HC1); (ii) at least three interleaved intrachromosomal SVs, four or more interchromosomal SVs, seven or more contiguous copy-number oscillations, and failure of the fragment-joins test (HC2); or (iii) a large number of rearrangements (≥40 intra- or interchromosomal SVs) accompanied by failure of the fragment-joins test (HC3). Chromosomes meeting a reduced set of criteria, defined by at least six interleaved intrachromosomal SVs, four to six adjacent copy-number oscillations between two states, failure of the fragment-joins test, and a significant chromosomal enrichment or exponential breakpoint distribution test, were classified as low-confidence chromothripsis events (LC). Unless otherwise specified, downstream analyses focused on both HC and LC chromothripsis events.

### Mutational signature analysis

Mutational signatures were identified using established computational frameworks^36^. Somatic mutation count matrices for SBS, small insertions and deletions (ID), and DBS were generated from final somatic variant call files using SigProfilerMatrixGenerator. Based on SBS96, ID83, and DBS78 mutational contexts, de novo mutational signature extraction was performed using SigProfilerExtractor^43^ with default parameters. Extracted signatures were subsequently matched to COSMIC reference signatures (v3.4) using the GRCh38 genome build. To estimate mutational signature activities at the individual tumor level, we applied the Mutational Signature Attribution (MSA) tool^44^, which uses a non-negative least squares (NNLS) algorithm with simulation-based optimization. MSA was used to fit the COSMIC signature set identified by SigProfilerExtractor to each tumor, with attribution thresholds automatically optimized by the pipeline. A conservative attribution strategy was applied using the no_CI_for_penalties = false option to determine optimal penalties. Pruned attributions were used for downstream analyses, in which 95% confidence intervals were computed for each signature and signature activities with a lower confidence bound of zero were excluded.

To minimize over-assignment of mutational signatures, analyses were stratified by tumor mutational burden, with high-mutational burden tumors (POLE and MSI subtypes) and low-mutational burden tumors (CN-High and CN-Low subtypes) analyzed separately. For indel and DBS signatures, which typically have lower mutation counts, mutational signature attributions from MSA were retained only if the reconstructed mutational profile showed an improvement of at least 5% in cosine similarity compared with the de novo extraction obtained using SigProfilerExtractor. Unless otherwise specified, a mutational signature was considered “present” in a tumor if at least 50 mutations were assigned to that signature and it contributed more than 5% of total mutations. For association analyses, signature presence was used for enrichment testing; alternatively, when a signature was present in more than 50% of tumors, signature activity was dichotomized based on values above or below the cohort median. These binary signature variables were subsequently used as dependent variables in multivariable logistic regression models.

### Copy-number and structural variant signature analysis

Copy-number alteration (CNA) and structural variant (SV) signatures were inferred *de novo* using SigProfilerExtractor, which applies a non-negative matrix factorization-based framework to identify recurrent genomic alteration patterns across tumors. CNA signatures were derived from CN48 profiles generated using SigProfilerMatrixGenerator, which encodes allele-specific copy-number segments according to absolute copy number, segment length, and allelic configuration while accounting for tumor ploidy. CNA segments were classified into three allelic states: heterozygous (copy number >0 on both alleles), loss of heterozygosity (copy number >0 on one allele and 0 on the other), and homozygous deletion (copy number 0 on both alleles). SV signatures were similarly inferred using SigProfilerExtractor based on SV32 profiles, which summarize structural variants by event type, size, and clustering properties. Tumor-level CN48 and SV32 feature matrices were decomposed to identify recurrent copy-number and structural rearrangement patterns across samples for downstream analyses, with COSMIC-defined classifications used for interpretative visualization and comparative analyses.

### Telomere length estimation

Telomere length was estimated from whole-genome sequencing data using TelSeq (v0.0.2)^45^. Reads containing at least seven consecutive TTAGGG or CCCTAA repeats were classified as telomeric. Read-group-level estimates were aggregated to generate sample-level telomere length values, with GC-content normalization applied using reads with 48-52% GC content.

### RNA-seq analysis

Normalized RNA-seq gene expression data for UCEC tumors were obtained from the TCGA PanCancer Atlas via cBioPortal, using RSEM-based quantification. Gene expression profiles were used to characterize transcriptional programs across molecular subtypes and to assess associations with genomic features. Pathway-level transcriptional activity was quantified using single-sample gene set enrichment analysis (ssGSEA) implemented in the GSVA R package. Curated gene sets were retrieved from the Molecular Signatures Database (MSigDB) using the msigdbr package, including the Hallmark and KEGG collections. ssGSEA scores were computed independently for each tumor to generate sample-level pathway activity profiles for downstream analyses. In parallel, expression levels of selected genes with established roles in proliferation, immune regulation, and DNA replication, including *MKI67*, *TOP2A*, and *CD274* (PD-L1), were analyzed at the gene level. Furthermore, tumor proliferation was assessed using a well-established, precomputed proliferation score obtained from the TCGA Pan-Cancer Atlas, reflecting tumor-intrinsic proliferative activity and applied consistently with its use in The Immune Landscape of Cancer study^46^. Gene expression differences across molecular subtypes and genomic strata were assessed using two-sided non-parametric Wilcoxon rank-sum tests. For analyses involving multiple gene sets or features, P values were adjusted for multiple testing using the Benjamini-Hochberg false discovery rate (FDR) procedure.

### Neoantigen analysis

Neoantigen load was obtained from the TCGA Pan-Cancer Atlas^46^. For each tumor, SNV-derived and indel-derived neoantigen burdens were analyzed separately and log_2_-transformed prior to downstream analyses. Differences in neoantigen load across UCEC molecular subtypes were assessed using two-sided non-parametric Wilcoxon rank-sum tests. For pairwise subtype comparisons, P values were adjusted for multiple testing using the Benjamini-Hochberg FDR procedure.

### Replication stress score estimation

Replication stress scores were derived from transcriptomic data using a previously published replication stress gene signature^47^. Gene expression values were standardized and weighted according to gene-specific coefficients obtained from a principal component-based model. Weighted expression values were summed to generate a single replication stress score per sample, which was subsequently Z-score normalized within each analysis cohort.

### Indel mutational signature analysis in CRISPR-Cas9 knockout models

To validate the association between mismatch repair (MMR) deficiency and the ID2-ID1 indel mutational signature imbalance, we reanalyzed whole-genome sequencing data from an independent CRISPR–Cas9–based isogenic cell line study^23^. This study used the immortalized human near-haploid HAP1 cell line to generate isogenic knockouts of DNA repair genes using CRISPR–Cas9. Somatic indel mutation catalogs were generated as ID83 matrices and subjected to de novo mutational signature extraction using SigProfiler-based pipelines, as described above. The similarity between extracted signatures and the reference ID2 signature was quantified using cosine similarity. To further characterize ID2- and ID1-associated indel classes, we quantified the frequencies of 1-bp thymine deletions (1:Del:T:5) and 1-bp thymine insertions (1:Ins:T:5) across different DNA repair knockout backgrounds.

### Statistical analysis

All statistical analyses were performed in R (v4.5.1; https://www.r-project.org/). Continuous variables were compared between groups using non-parametric Mann-Whitney (Wilcoxon rank-sum) tests. Associations between categorical variables were evaluated using Fisher’s exact test or logistic regression, with adjustment for relevant covariates when appropriate. Survival analyses were performed using Kaplan-Meier estimates with differences between groups assessed by two-sided log-rank tests and using Cox proportional hazards regression models to evaluate associations with overall survival and progression-free interval. Cox models were adjusted for relevant clinical covariates, including age at diagnosis, tumor stage, grade and histological or molecular subtype, as applicable. All statistical tests were two-sided, and P values < 0.05 were considered statistically significant. For analyses involving multiple comparisons, P values were adjusted using the Benjamini-Hochberg FDR method, and adjusted P values are reported.

## Data availability

Access to TCGA controlled data can be requested through dbGaP (study accession: phs000178.v11.p8). Raw WGS data (BAM files) for the TCGA UCEC cohort were obtained from the Genomic Data Commons (GDC) data portal (https://portal.gdc.cancer.gov/) under standard TCGA data access policies. Processed RNA-seq data (quantified as RSEM values) for the TCGA UCEC cohort are available via cBioPortal (https://www.cbioportal.org/), using the study name “Uterine Corpus Endometrial Carcinoma (TCGA, PanCancer Atlas)”.

## Code availability

The bioinformatics pipelines used for whole-genome sequencing analysis are available at https://github.com/xtmgah/Sherlock-Lung. The source code used to generate the main figures presented in this manuscript is accessible at https://github.com/xtmgah/TCGA-WGS-Mansucripts.

## Acknowledgments

This work utilized the computational resources of the NIH HPC Biowulf cluster (https://hpc.nih.gov).

This research was supported by the Intramural Research Program of the National Institutes of Health (NIH). The contributions of the NIH author(s) were made as part of their official duties as NIH federal employees, are in compliance with agency policy requirements, and are considered Works of the United States Government. However, the findings and conclusions presented in this paper are those of the author(s) and do not necessarily reflect the views of the NIH or the U.S. Department of Health and Human Services.

This project has been funded in whole or in part with Federal funds from the National Cancer Institute, National Institutes of Health, under Contract No. 75N91019D00024. The content of this publication does not necessarily reflect the views or policies of the Department of Health and Human Services, nor does mention of trade names, commercial products, or organizations imply endorsement by the U.S. Government.

## Author Contributions

Conceptualization, SJC, TZ; Methodology, JS, MZ, JL, VB, TZ; Formal Analysis, JS, MZ, TZ; Resources, BZ, GW, VB, SJC, TZ; Data Curation, JS, MZ, TV, YK, SC, WZ, WL, AMM, TZ; Writing – Original Draft, JS, TZ; Writing – Review & Editing, JS, MZ, JL, GW, BZ, VB, SJC, TZ; Visualization, JS, MZ, TZ; Supervision, TZ.

## Declaration of Interests

All authors declare that they have no competing interests.

### Inclusion & Ethics Statement

This study utilized publicly available genomic, and transcriptomic data from The Cancer Genome Atlas (TCGA), ensuring broad transparency and equitable access to resources. No new human subjects or biological samples were collected. All analyses were performed in compliance with the TCGA data usage policies and NIH ethical guidelines.

We acknowledge the contributions of the diverse TCGA participant cohort and the researchers who generated and curated these datasets. Authorship reflects contributions across scientific roles including data analysis, interpretation, and manuscript preparation.

---

**Extended Data Fig. 1:**
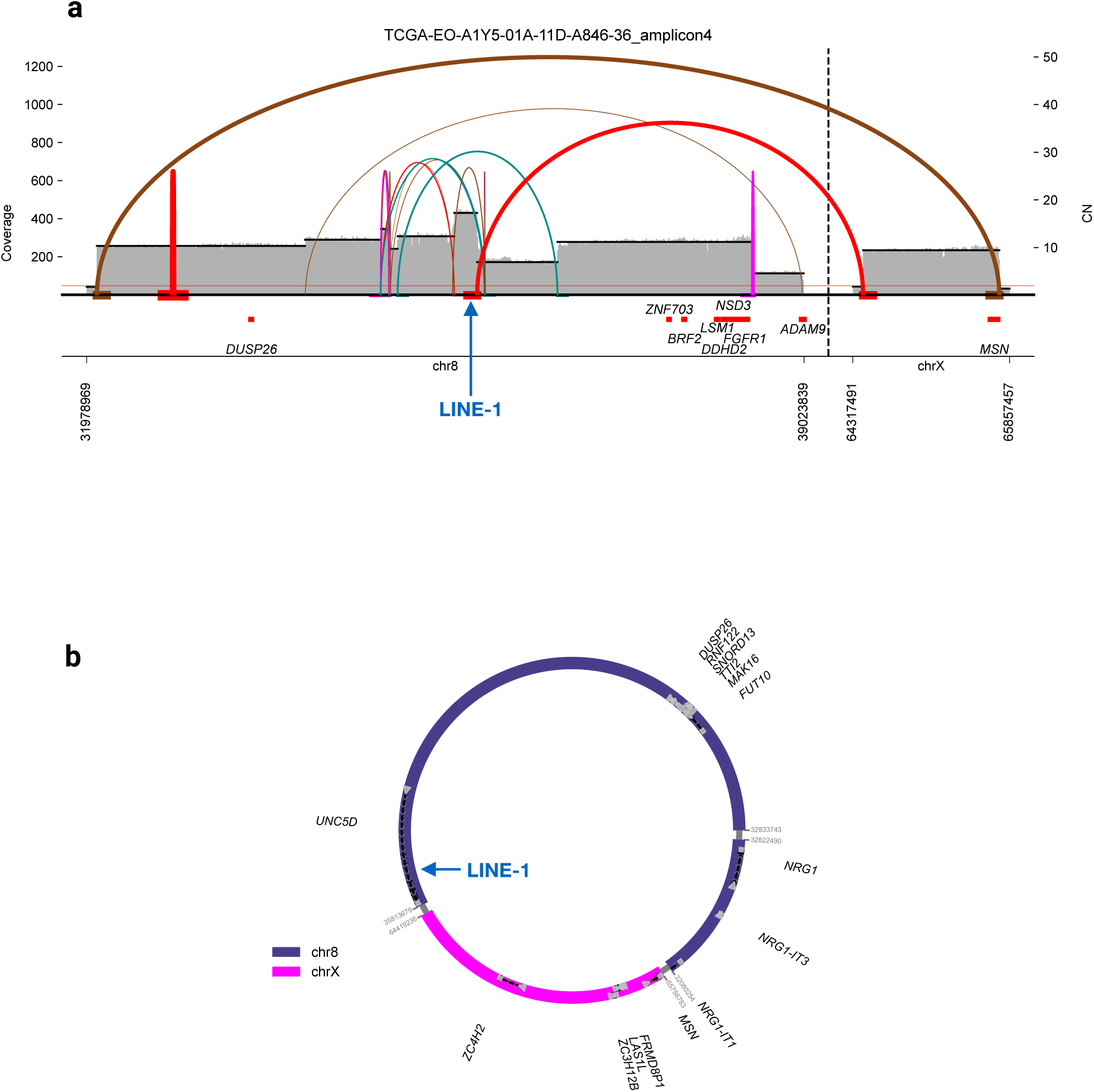
Association between LINE-1 insertions and focal amplification mechanisms a, Representative example of a LINE-1 insertion detected within an ecDNA structure. Coverage (left y-axis) and copy number (CN; right y-axis) profiles across chromosome 2 are shown for tumor sample *TCGA-EO-A1Y5-01A*. Black segments indicate segmented copy number states, with focal high-level amplification consistent with ecDNA. Curved arcs represent inferred rearrangement junctions defining the ecDNA structure. Vertical colored lines indicate structural variant breakpoints, and the position of the LINE-1 insertion is highlighted by a blue arrow. Gene annotations are shown along the genomic coordinate axis. **b,** Circular representation of the constructed ecDNA structure shown in **a**, illustrating the genomic segments involved in the ecDNA amplicon and their rearranged order. Segments originating from chromosome 2 (brown) and chromosome 20 (light blue) are indicated. The position of the LINE-1 insertion within the ecDNA is marked by a blue arrow. Selected genes located on the ecDNA are annotated around the circle.

**Extended Data Fig. 2:**
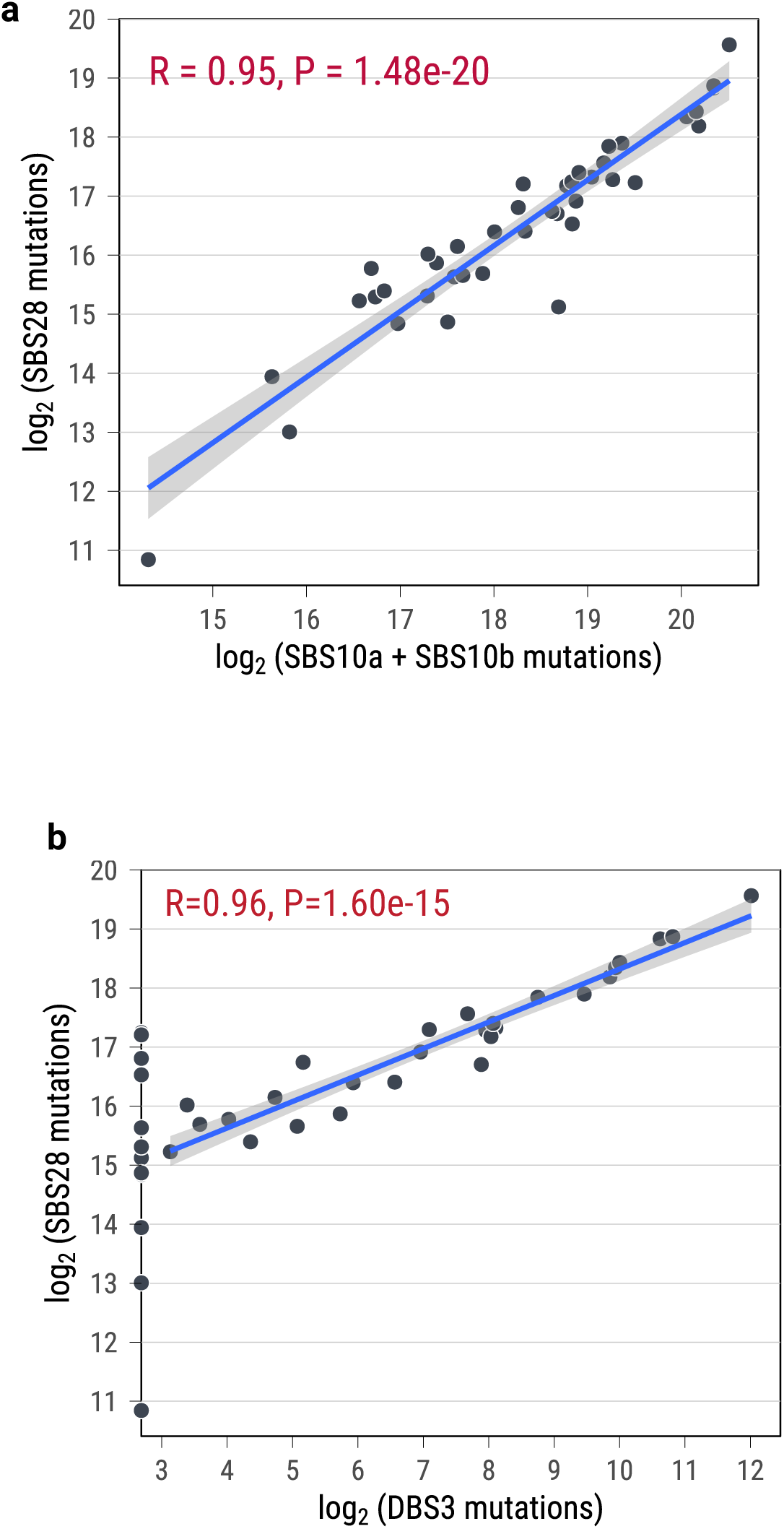
Pearson correlation between SBS28 and *POLE* exonuclease deficiency-associated mutational signatures a, Scatter plot showing the relationship between SBS28 mutation counts and the combined mutation counts of SBS10a and SBS10b across tumors. **b,** Scatter plot showing the relationship between SBS28 mutation counts and DBS3 mutation counts across tumors. Mutation counts are log_₂_-transformed. The solid line indicates the linear regression fit, with the shaded area representing the 95% confidence interval. The Pearson correlation coefficient (R) and two-sided P value are shown.

**Extended Data Fig. 3:**
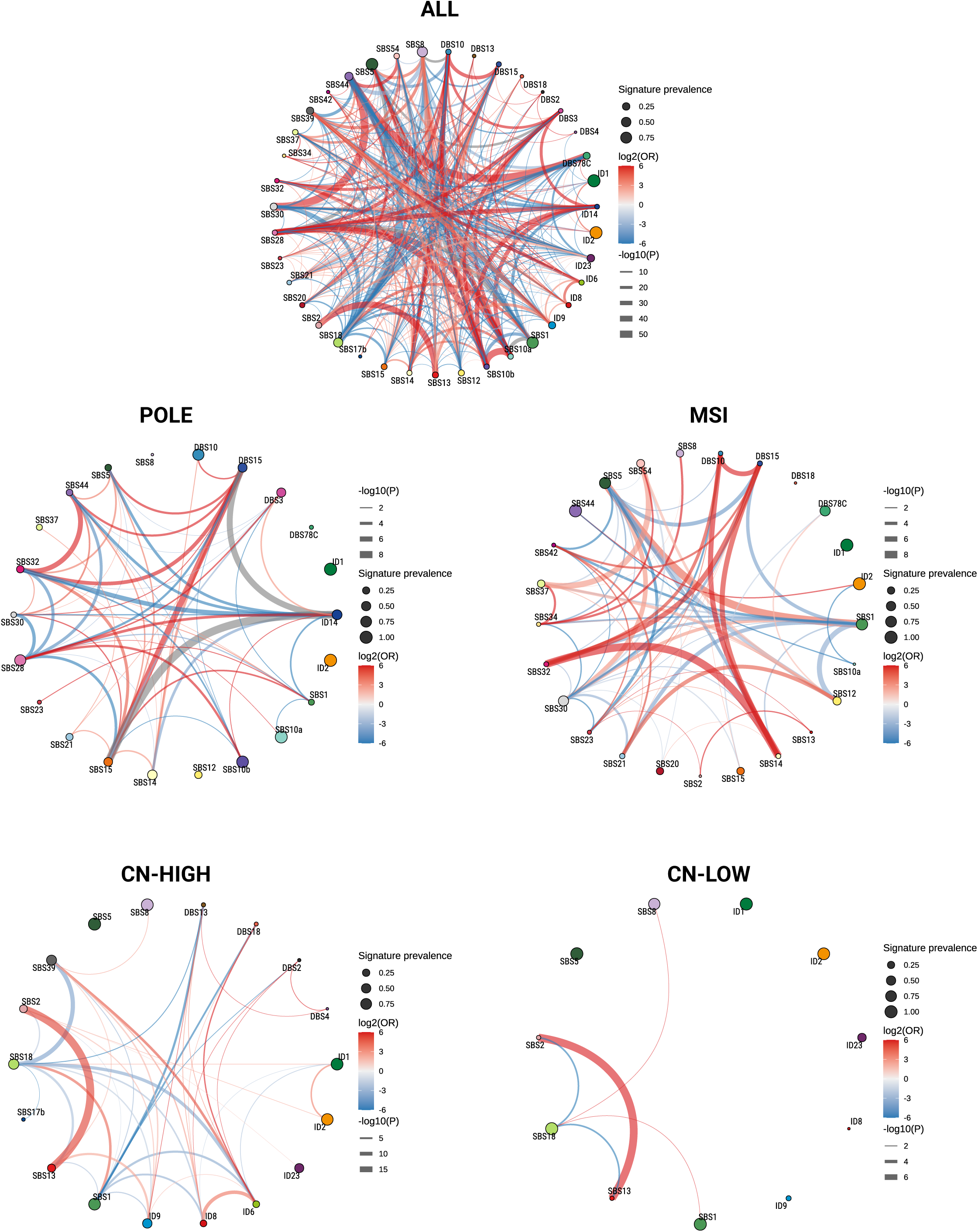
Pairwise association networks of mutational signatures across UCEC molecular subtypes. Circular network plots depict pairwise associations between mutational signatures across all tumors (ALL) and stratified by molecular subtype (POLE, MSI, CN-High, and CN-Low). Nodes represent individual COSMIC mutational signatures, with node size proportional to the prevalence of each signature within the corresponding group. Edges indicate statistically significant associations between signatures, with edge color denoting the direction and magnitude of association (log_₂_ odds ratio; red, positive association; blue, negative association) and edge width proportional to statistical significance (−log_₁₀_ P value). These networks reveal subtype-specific patterns of co-occurrence (red) and mutual exclusivity (blue) among mutational processes, including negative associations between SBS28 and MSI-related signatures in POLE tumors.

**Extended Data Fig. 4:**
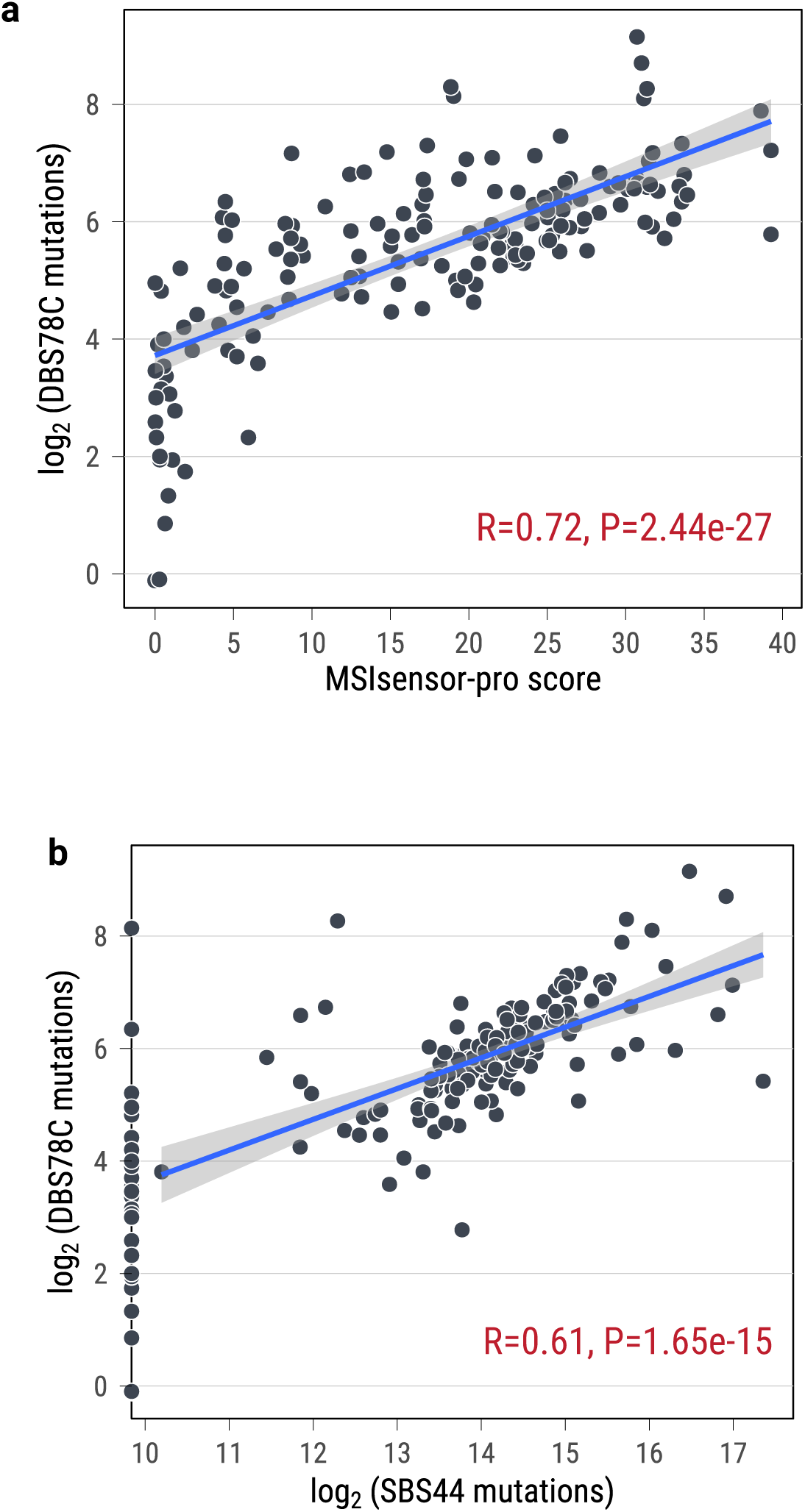
Pearson correlations between DBS78C mutations and MSI-associated features a, Scatter plot showing the relationship between DBS78C mutation counts and microsatellite instability (MSI) scores estimated by MSIsensor-pro. **b,** Scatter plot showing the relationship between DBS78C mutation counts and SBS44 mutation counts across tumors. Mutation counts are log_₂_-transformed. The solid line indicates the linear regression fit, with the shaded region representing the 95% confidence interval. The Pearson correlation coefficient (R) and two-sided P value are shown.

**Extended Data Fig. 5:**
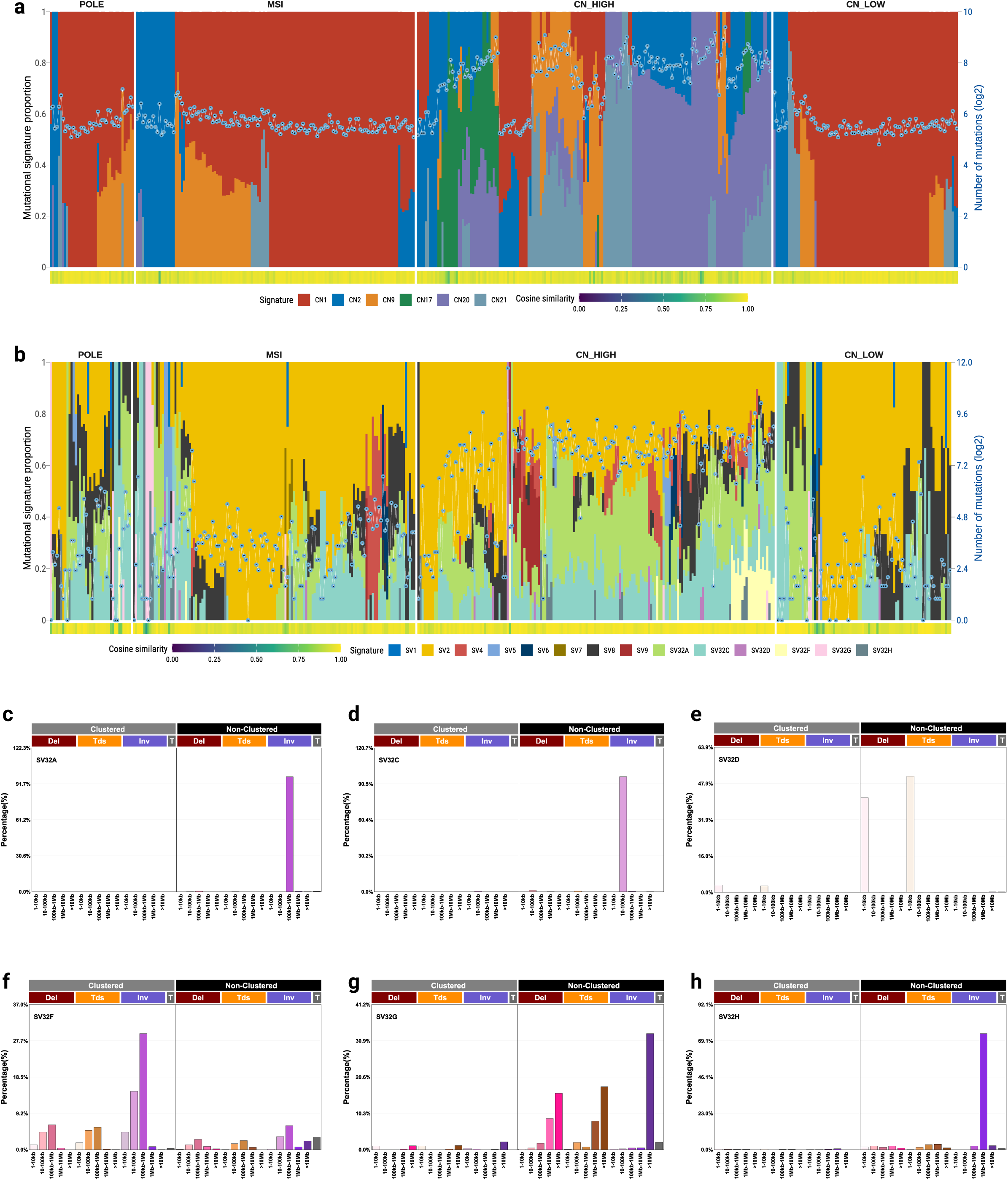
Copy-number and structural variant signature landscapes across UCEC molecular subtypes. **a**, Sample-level profiles of copy-number (CN) signatures across UCEC tumors, ordered by molecular subtype (POLE, MSI, CN-High, and CN-Low). Stacked bars indicate the relative contribution of individual CN signatures within each tumor. Dots denote the total number of copy-number events per sample (log_₂_-transformed). The color bar below indicates cosine similarity between original and reconstructed profiles. **b**, Sample-level profiles of structural variant (SV) signatures across UCEC tumors, ordered by molecular subtype. Stacked bars represent the relative contribution of individual SV signatures within each tumor. Dots indicate the total number of SV events per sample (log_₂_-transformed). The color bar below indicates cosine similarity. **c-h,** *De novo* structural variant (SV) signature profiles (SV32A, SV32C, SV32D, SV32F, SV32G, and SV32H) identified in UCEC tumors. Bar plots show the relative contributions of deletions (Del), tandem duplications (Tds), inversions (Inv), and translocations (T) across size categories.

**Extended Data Fig. 6:**
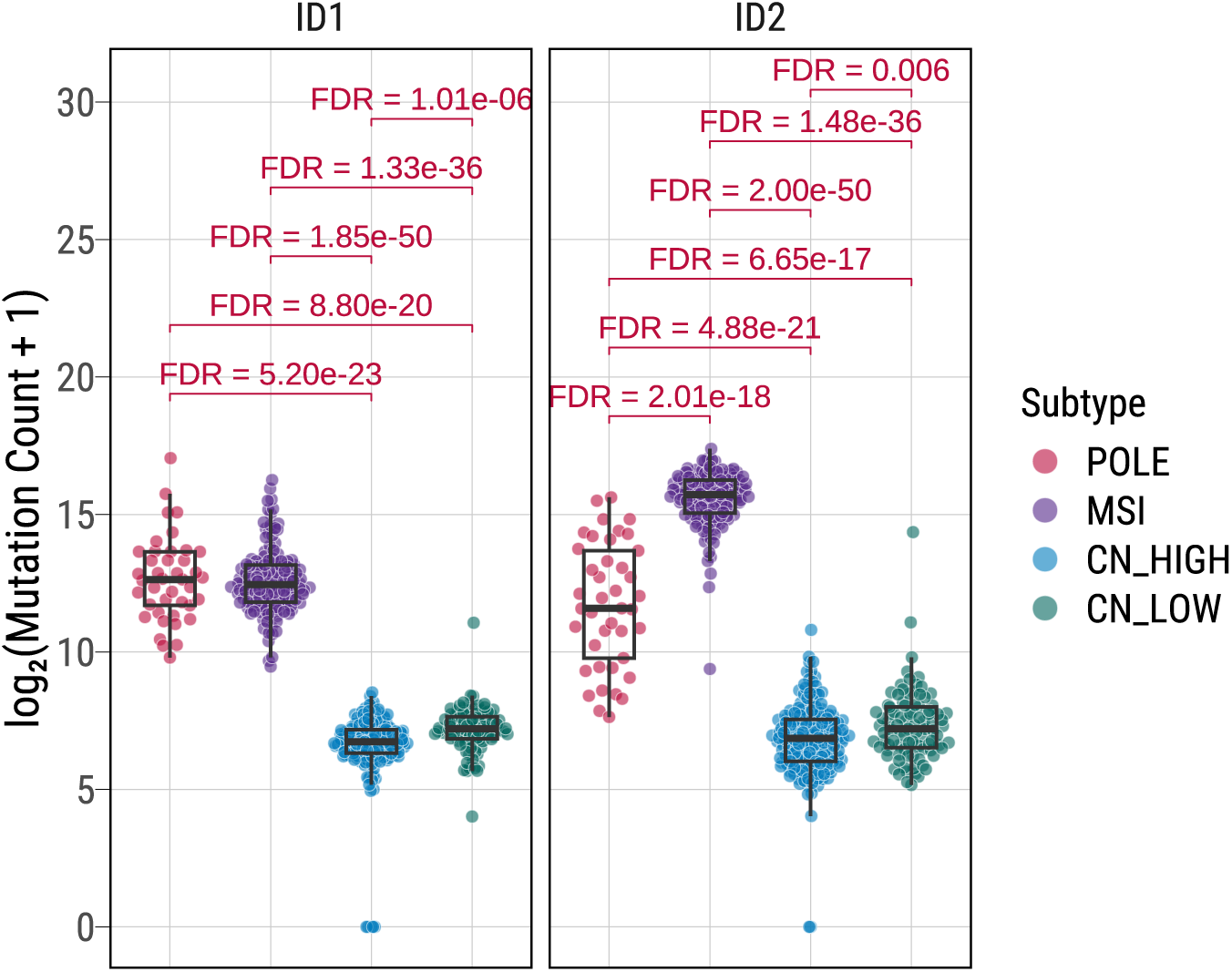
Distributions of ID1 and ID2 mutation burdens across UCEC molecular subtypes. Boxplots showing the distribution of mutations attributed to signatures ID1 (left) and ID2 (right) across UCEC molecular subtypes. Mutation counts are log_₂_-transformed. Each dot represents an individual tumor. Boxes indicate the median and interquartile range, with whiskers extending to 1.5× the interquartile range. Pairwise statistical comparisons between subtypes were performed using a two-sided Wilcoxon rank-sum test with FDR correction, and adjusted FDR values are shown.

**Extended Data Fig. 7:**
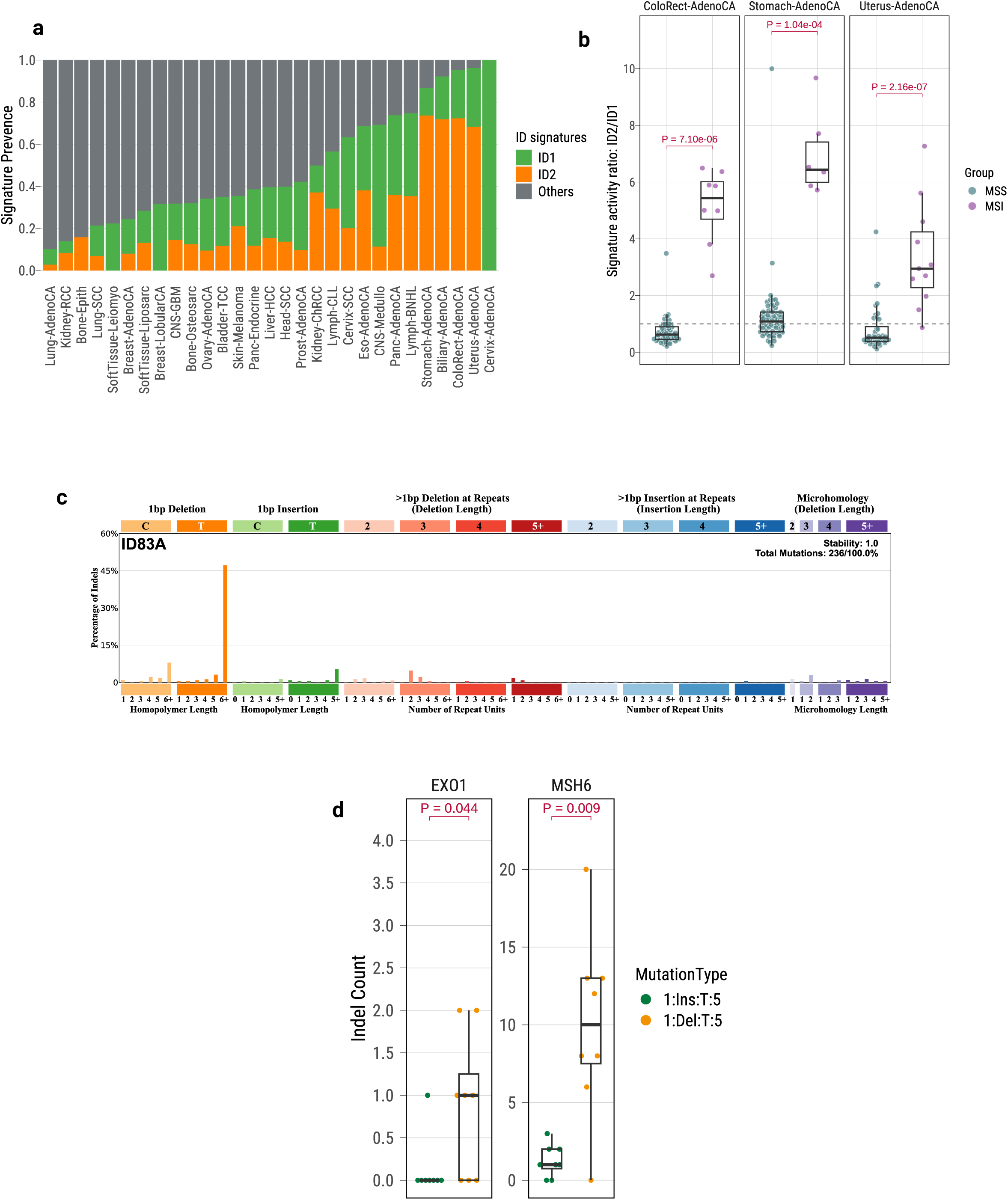
Validation of ID1 and ID2 signature imbalance across pan-cancer datasets and DNA repair-deficient models a, Stacked bar plot showing the prevalence of indel mutational signatures ID1 and ID2 across multiple tumor types from the PCAWG whole-genome sequencing dataset. Bars indicate the relative contributions of ID1, ID2, and other indel signatures within each cancer type. **b,** Boxplots showing the ratio of ID2 to ID1 signature activity in colorectal, stomach, and uterine adenocarcinomas from the PCAWG dataset, stratified by microsatellite status (microsatellite stable, MSS; microsatellite instability, MSI). Each point represents an individual tumor. Statistical comparisons were performed using two-sided Wilcoxon rank-sum tests, with P values shown. **c,** Indel mutational profile of the ID83A signature derived from CRISPR-Cas9-based knockout experiments of DNA repair genes, illustrating the distribution of 1-bp insertions and deletions, indels at repeat sequences, and microhomology-mediated deletions. **d,** Boxplots showing counts of 1-bp thymine insertions (ID1-type) and 1-bp thymine deletions (ID2-type) in CRISPR-Cas9-based knockout models of *EXO1* and *MSH6*. Each point represents an individual sample. Statistical comparisons between mutation types were performed using two-sided Wilcoxon rank-sum tests, with P values shown.

**Extended Data Fig. 8:**
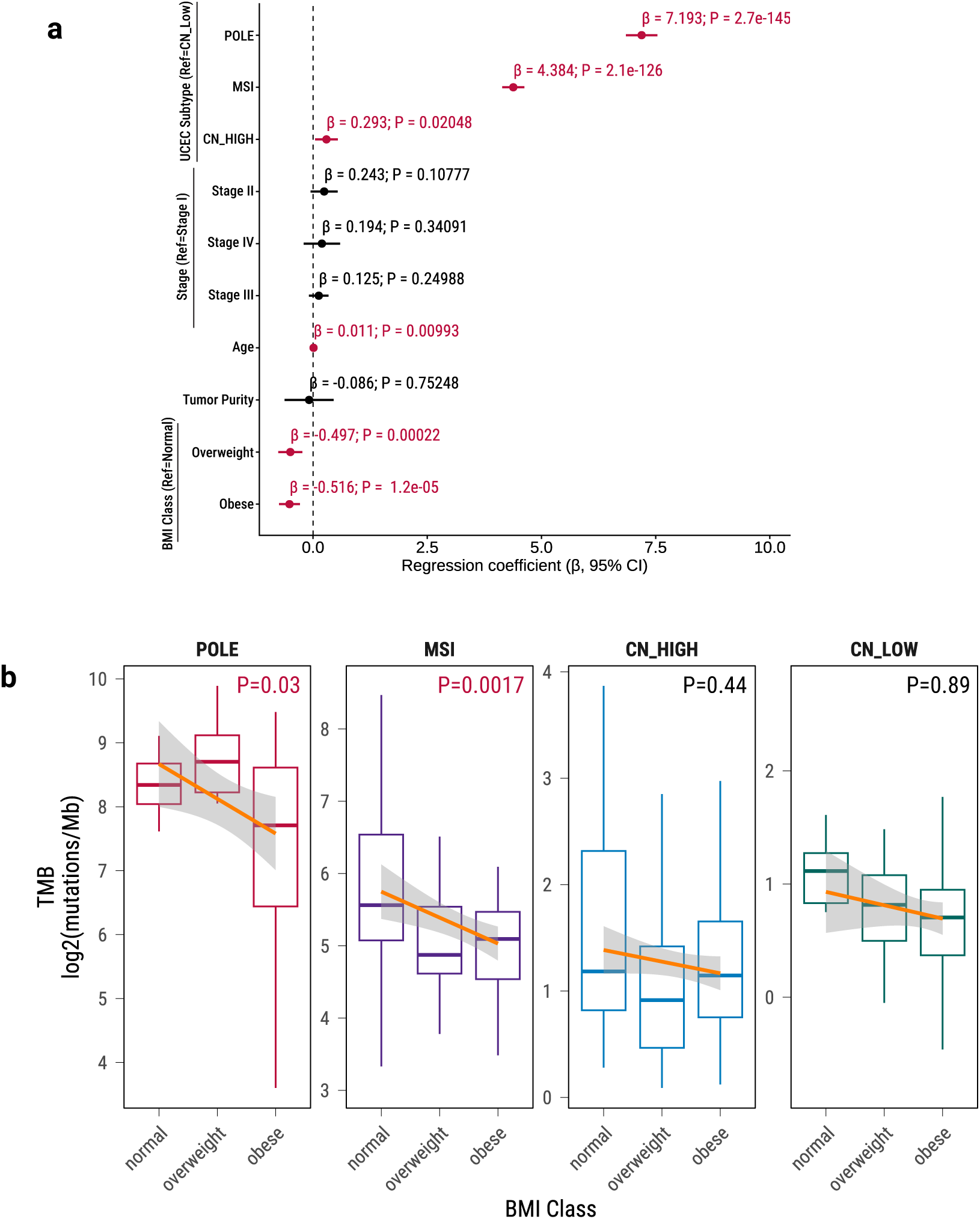
Association between body mass index and tumor mutational burden in UCEC a, Multivariable regression analysis assessing associations with tumor mutational burden (TMB) in TCGA-UCEC. Variables include UCEC molecular subtype (reference = CN-Low), age at diagnosis, tumor purity, tumor stage (reference = stage I), and BMI class (reference = normal). Points represent regression coefficients (β), and horizontal lines indicate 95% confidence intervals. Two-sided P values are shown. **b,** Association between BMI class (normal, overweight, obese) and tumor mutational burden within each UCEC molecular subtype. Boxplots show log_₂_-transformed TMB (mutations per megabase). Solid lines indicate fitted trends across BMI classes within each subtype, with shaded regions representing 95% confidence intervals. P values indicate trend significance.

**Supplementary Fig. 1: Sequencing depth, tumor purity, histological composition, and age distribution across UCEC molecular subtypes**

**a,** Distribution of whole-genome sequencing depth for tumor samples and matched blood or normal samples across the UCEC cohort. Density plots show the sequencing depth distributions, with dashed vertical lines indicating median value for each sample type.

**b,** Distribution of tumor purity across UCEC tumors estimated from whole-genome sequencing data. The density plot and accompanying boxplot summarize the overall distribution, with the median tumor purity indicated.

**c,** Histological composition of UCEC tumors across molecular subtypes. Stacked bar plots show the number of tumors within each subtype stratified by primary diagnosis, as annotated in TCGA clinical records.

**d,** Age at diagnosis across UCEC molecular subtypes. Each dot represents an individual patient. Boxplots summarize the distribution within each subtype, with the median and interquartile range shown. FDR-adjusted P values for pairwise comparisons are indicated.

**Supplementary Fig. 2: Clonal copy-number classification across UCEC tumors**

Genome-wide copy-number alteration (CNA) profiles across UCEC tumors, hierarchically clustered based on relative copy-number patterns. Heatmap colors represent relative copy-number states, ranging from loss to high-level amplification, across autosomes, with tumors shown in rows and genomic segments ordered by chromosomal position along the x-axis. Tumors are grouped into four major copy-number clusters (C1-C4), as indicated by the dendrogram. The top panel shows the aggregate frequency of copy-number gains and losses across the cohort. Side annotations indicate tumor purity, WGD status, and UCEC molecular subtype.

**Supplementary Fig. 3: Genomic distribution of germline LINE-1 source elements across UCEC molecular subtypes**

Genome-wide distribution of germline LINE-1 source elements giving rise to somatic LINE-1 transductions across UCEC molecular subtypes. For each subtype, vertical bars indicate the average transduction source frequency along each chromosome, plotted by genomic position. Chromosomes are shown along the x-axis, and the y-axis indicates the average transduction source frequency. Recurrent germline LINE-1 source loci are labeled. The overall distribution pattern of dominant germline source elements is consistent across molecular subtypes.

**Supplementary Fig. 4: Associations between focal amplification mechanisms, copy-number clusters, and whole-genome doubling in CN-High tumors**

**a-d**, Proportions of CN-High tumors harboring different focal amplification architectures, including (**a**) linear amplification, (**b**) ecDNA, (**c**) BFB, and (**d**) complex non-cyclic amplification, stratified by copy-number cluster (C1 versus C3). Stacked bar plots show the percentage of tumors with (TRUE) or without (FALSE) the indicated amplification type within each cluster. Odds ratios (ORs) and P values are shown for each comparison.

**e-h**, Proportions of CN-High tumors harboring different focal amplification architectures, including (**e**) linear amplification, (**f**) ecDNA, (**g**) BFB, and (**h**) complex non-cyclic amplification, stratified by WGD status (WGD versus non-WGD). Stacked bar plots show the percentage of tumors with (TRUE) or without (FALSE) the indicated amplification type within each group. Odds ratios (ORs) and P values are shown for each comparison.

**Supplementary Fig. 5: Expression of amplified oncogenes stratified by focal amplification mechanisms**

mRNA expression levels of selected oncogenes frequently involved in focal amplifications in UCEC tumors. Expression values are shown as log_2_-transformed RNA-seq RSEM values. Each dot represents an individual tumor. Tumors harboring focal amplifications are highlighted and colored by amplification architecture, including linear amplification, ecDNA, BFB, and complex non-cyclic amplification, while gray points indicate tumors without focal amplification of the corresponding gene. Boxplots summarize the distribution of expression levels across all tumors, with the median and interquartile range indicated.

**Supplementary Fig. 6: Single-sample gene set enrichment analysis of ecDNA-positive and ecDNA-negative tumors**

Single-sample gene set enrichment analysis (ssGSEA) comparing ecDNA-positive and ecDNA-negative tumors across all UCEC tumors (**a,b**) and within the CN-High subtype (**c,d**). Each point represents a gene set, plotted by the difference in ssGSEA scores between ecDNA-positive and ecDNA-negative tumors (ecDNA − non-ecDNA). **a,** Hallmark gene sets across all tumors. **b,** KEGG gene sets across all tumors. **c,** Hallmark gene sets within CN-High tumors. **d,** KEGG gene sets within CN-High tumors.

**Supplementary Fig. 7: Survival analyses stratified by ecDNA status**

**a-b**, Kaplan-Meier curves of overall survival (**a**) and progression-free interval (**b**) stratified by ecDNA status.

**c-d**, Kaplan-Meier curves of overall survival (**c**) and progression-free interval (**d**) comparing ecDNA-positive and ecDNA-negative tumors within the CN-High subtype.

**e-f**, Kaplan-Meier curves of overall survival (**e**) and progression-free interval (**f**) for all tumors stratified by ecDNA-associated gene categories, including *MYC* ecDNA, *ERBB2* ecDNA, other ecDNA, and non-ecDNA tumors. Log-rank test P values are indicated in each panel.

The x-axis represents overall survival time (days) for overall survival plots and progression-free interval time (days) for PFI plots.

**Supplementary Fig. 8: Association between ecDNA and chromothripsis within the CN-High subtype**

Stacked bar plots showing the proportion of tumors with and without chromothripsis stratified by ecDNA status among CN-High UCEC tumors. Numbers within bars indicate the number of samples and corresponding percentages in each category. The association between ecDNA status and chromothripsis was assessed using Fisher’s exact test, with the odds ratio (OR) and P value shown above the plot.

**Supplementary Fig. 9: Survival analyses stratified by chromothripsis**

Kaplan-Meier curves of overall survival (**a**) and progression-free interval (**b**) stratified by combined CN-High and chromothripsis status. Tumors were grouped as CN-High with chromothripsis (CN-High/chromothripsis+), CN-High without chromothripsis (CN-High/chromothripsis-), and all remaining tumors (Others). Log-rank test P values are shown in each panel. Tick marks indicate censored observations. The x-axis represents overall survival time (days) for overall survival plots and progression-free interval time (days) for PFI plots.

**Supplementary Fig. 10: LINE-1 insertion burden stratified by chromothripsis status within the CN-High subtype**

Boxplots showing the distribution of LINE-1 insertion burden, quantified as log2(TE count + 1), in CN-High tumors with and without chromothripsis. Each dot represents an individual tumor. Boxes indicate the median and interquartile range, with whiskers extending to 1.5× the interquartile range. Statistical significance was assessed using a two-sided Wilcoxon rank-sum test, with the P value indicated.

**Supplementary Fig. 11: Comparison of indel counts attributed to each ID mutational signature between CN-High and CN-Low tumors**

Boxplots showing the distribution of indel mutational signature counts in CN-High and CN-Low tumors. Each dot represents an individual tumor. Boxes indicate the median and interquartile range, with whiskers extending to 1.5× the interquartile range. Statistical comparisons between CN-High and CN-Low tumors were performed using a two-sided Wilcoxon rank-sum test, with P values indicated above each panel.

**Supplementary Fig. 12: Distributions of mutational signature burdens across UCEC molecular subtypes**

Scatter plots showing the distributions of mutation counts for all SBS, DBS, and ID signatures across individual tumors, colored by molecular subtype. Mutation counts are displayed on a log_₁₀_ scale. Each dot represents an individual tumor, and horizontal lines indicate the median value for each signature. Numbers above the plot indicate the total number of tumors contributing data to each signature.

**Supplementary Fig. 13: Replication timing profiles of ID1 mutations across UCEC molecular subtypes**

Bar plots showing the normalized density of ID1 small insertion and deletion mutations across replication timing bins, ordered from early- to late-replicating regions, stratified by UCEC molecular subtype. Dashed lines indicate smoothed trends across replication timing bins. Replication timing is shown from early to late along the x-axis.

**Supplementary Fig. 14: Expression of proliferation and immune checkpoint markers across UCEC molecular subtypes**

Boxplots showing mRNA expression levels of *MKI67*, *TOP2A*, and *CD274* across UCEC molecular subtypes. Expression values are shown as log₂(RNA-seq RSEM+1). Each dot represents an individual tumor. Boxes indicate the median and interquartile range, with whiskers extending to 1.5× the interquartile range. Pairwise statistical comparisons between subtypes were performed using a two-sided Wilcoxon rank-sum test with FDR correction, and FDR adjusted P values are shown.

**Supplementary Fig. 15: Telomere length comparison across UCEC molecular subtypes**

Boxplots showing telomere length ratios, calculated as log_₂_(tumor/normal), across UCEC molecular subtypes. Each dot represents an individual tumor. Boxes indicate the median and interquartile range, with whiskers extending to 1.5× the interquartile range. Pairwise statistical comparisons between subtypes were performed using a two-sided Wilcoxon rank-sum test with FDR correction, and FDR adjusted P values are shown.

**Supplementary Fig. 16: Detection frequency of IIV31 across UCEC molecular subtypes**

Stacked bar plots showing the proportion of tumors with and without detectable IIV31 across UCEC molecular subtypes. Numbers within bars indicate the number of tumors and corresponding percentages in each category. Differences in IIV31 detection frequency between subtypes were assessed using Fisher’s exact test, with the OR and P value indicated.

**Supplementary Fig. 17: Distribution of body mass index categories across UCEC molecular subtypes**

Stacked bar plots showing the distribution of BMI categories (normal, overweight, and obese) across UCEC molecular subtypes. BMI categories were defined according to standard clinical cutoffs (normal, BMI < 25 kg/m²; overweight, BMI 25-29.9 kg/m²; obese, BMI ≥ 30 kg/m²). Numbers within bars indicate the number of patients and corresponding percentages in each category.

**Supplementary Data 1: Demographic, clinical, and tumor characteristics of 440 TCGA UCEC patients.**

**Supplementary Data 2: Comprehensive landscape of mobile element insertions (MEIs) across TCGA UCEC tumor genomes.**

**Supplementary Data 3: Classification and burden of focal amplification mechanisms in TCGA UCEC tumors.**

**Supplementary Data 4: Annotation of chromothripsis events and associated structural variation features in TCGA UCEC tumors.**

**Supplementary Data 5: Activities of single base substitution (SBS), doublet base substitution (DBS), and small insertion/deletion (ID) mutational signatures in TCGA UCEC tumors.**

**Supplementary Data 6: Mutational profile of the newly identified doublet base substitution (DBS) signature DBS78C in TCGA UCEC tumors.**

**Supplementary Data 7: Structural variation–derived mutational signature activities in TCGA UCEC tumors.**

**Supplementary Data 8: Copy number variation–derived mutational signature activities in TCGA UCEC tumors.**

