## Supplementary Figures for "Endogenous mutational mechanisms and metabolic context shape endometrial cancer"

#### Supplementary Fig. 1

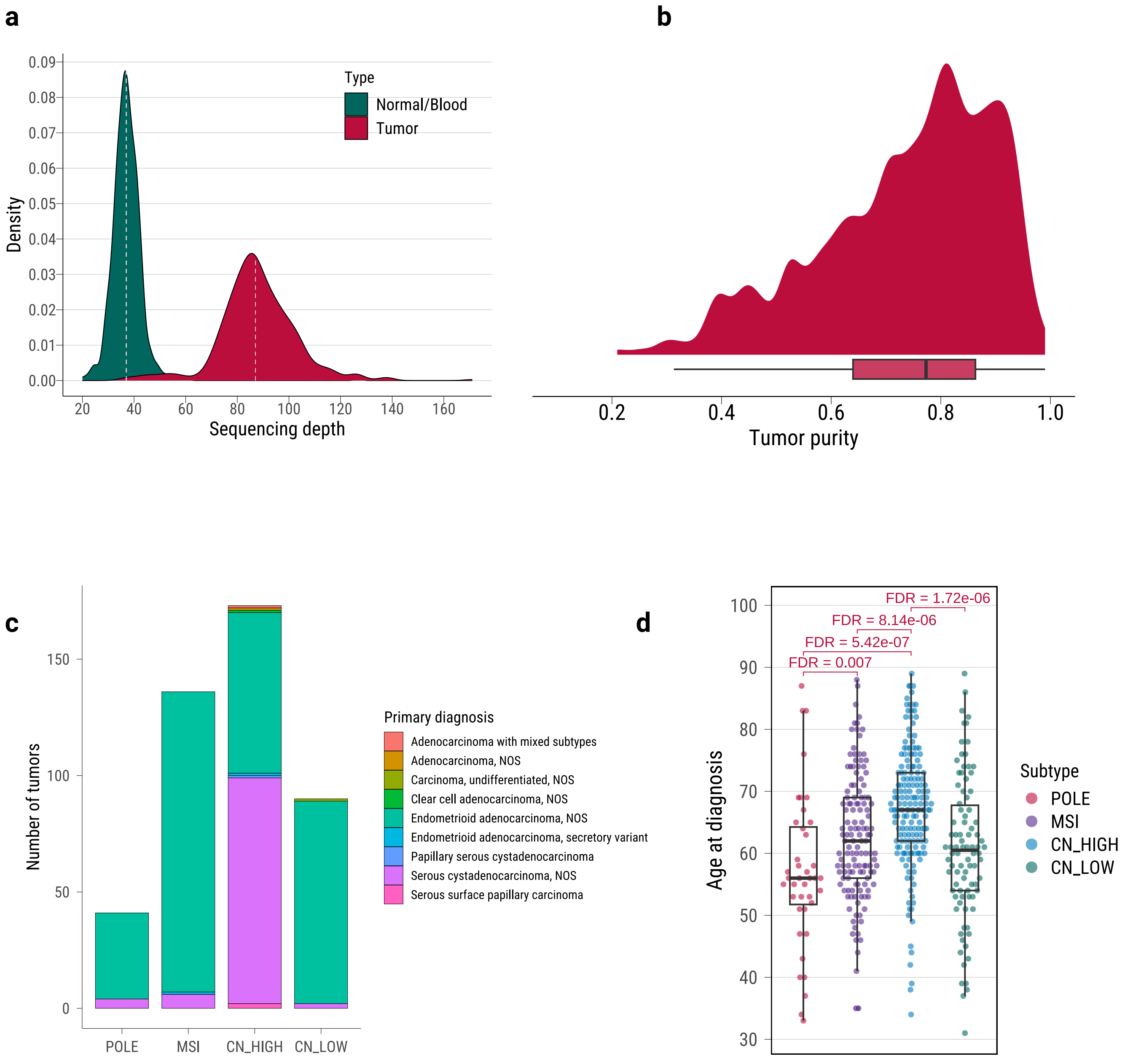

### Supplementary Fig. 2

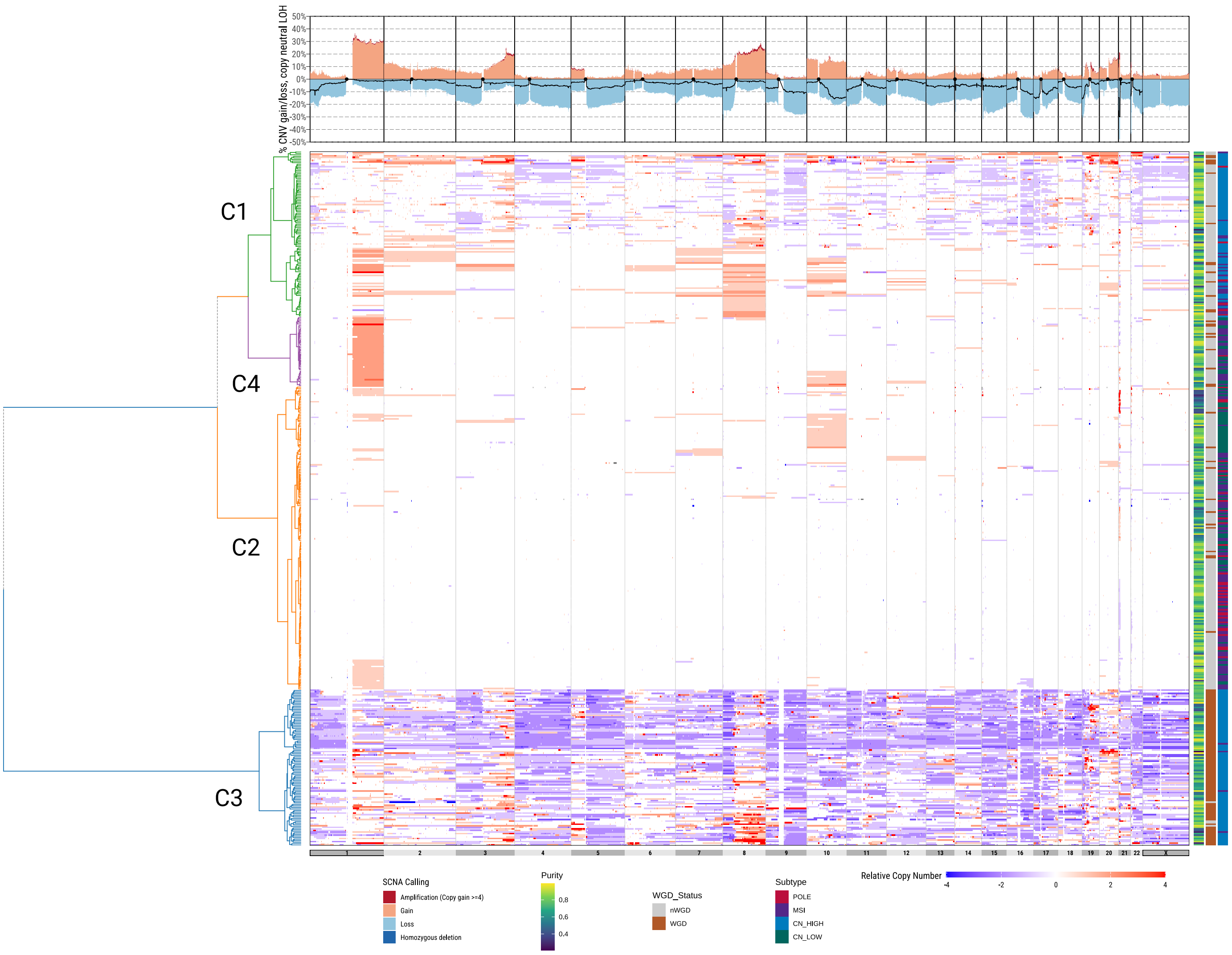

### Supplementary Fig. 3

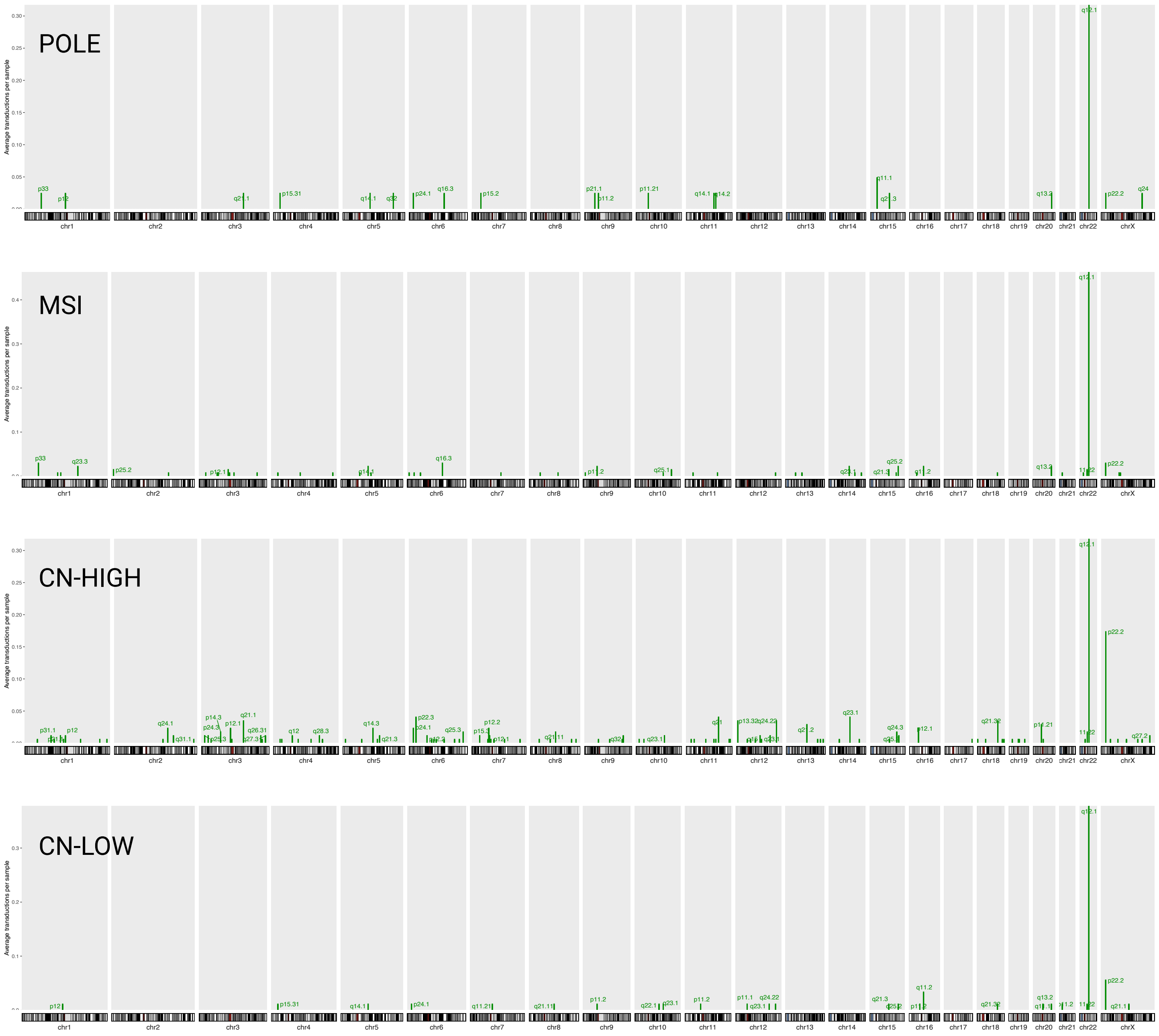

**Supplementary Fig. 4**

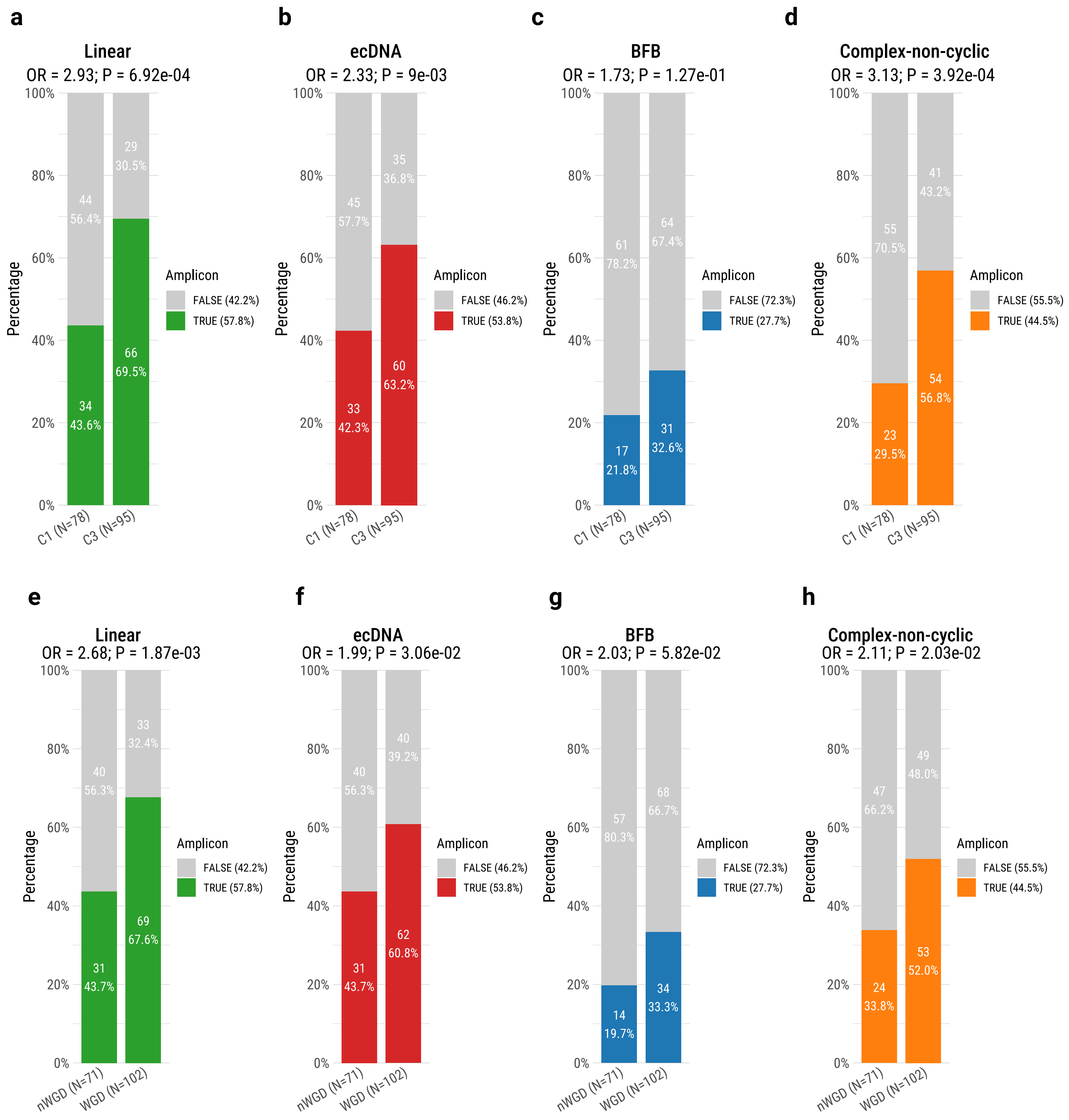

**Supplementary Fig. 5**

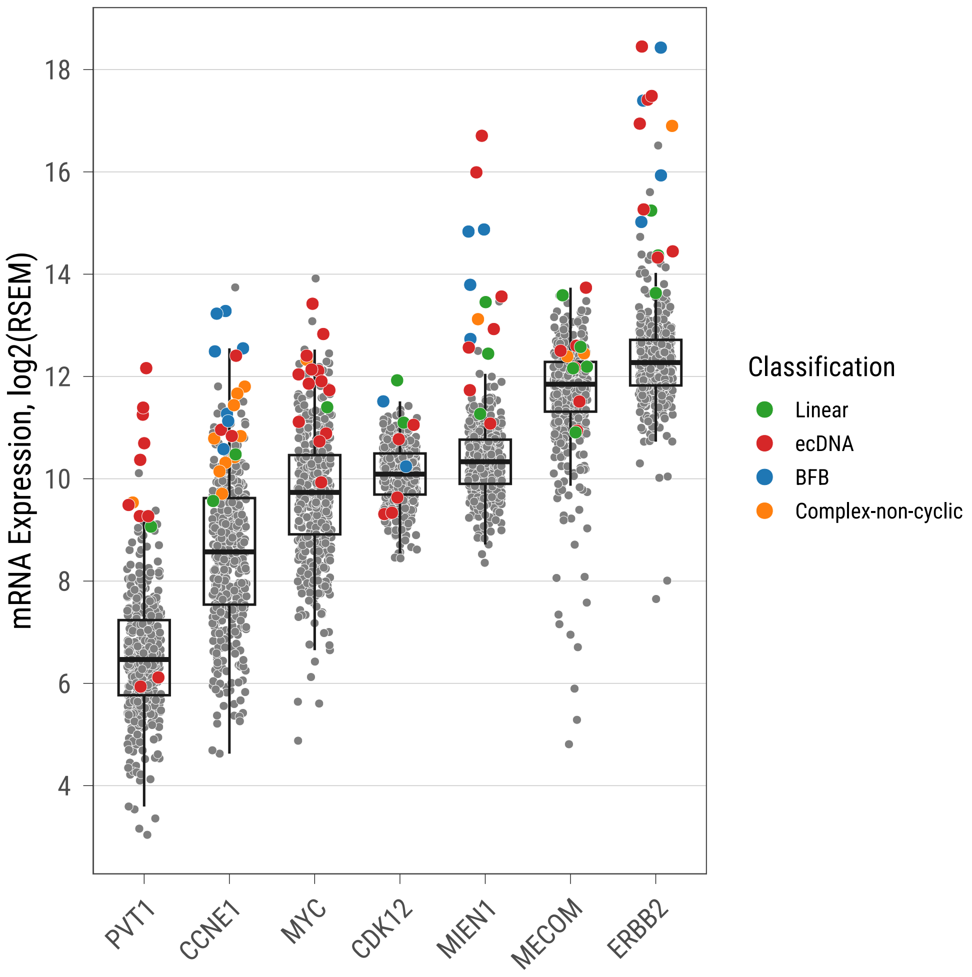

#### Supplementary Fig. 6

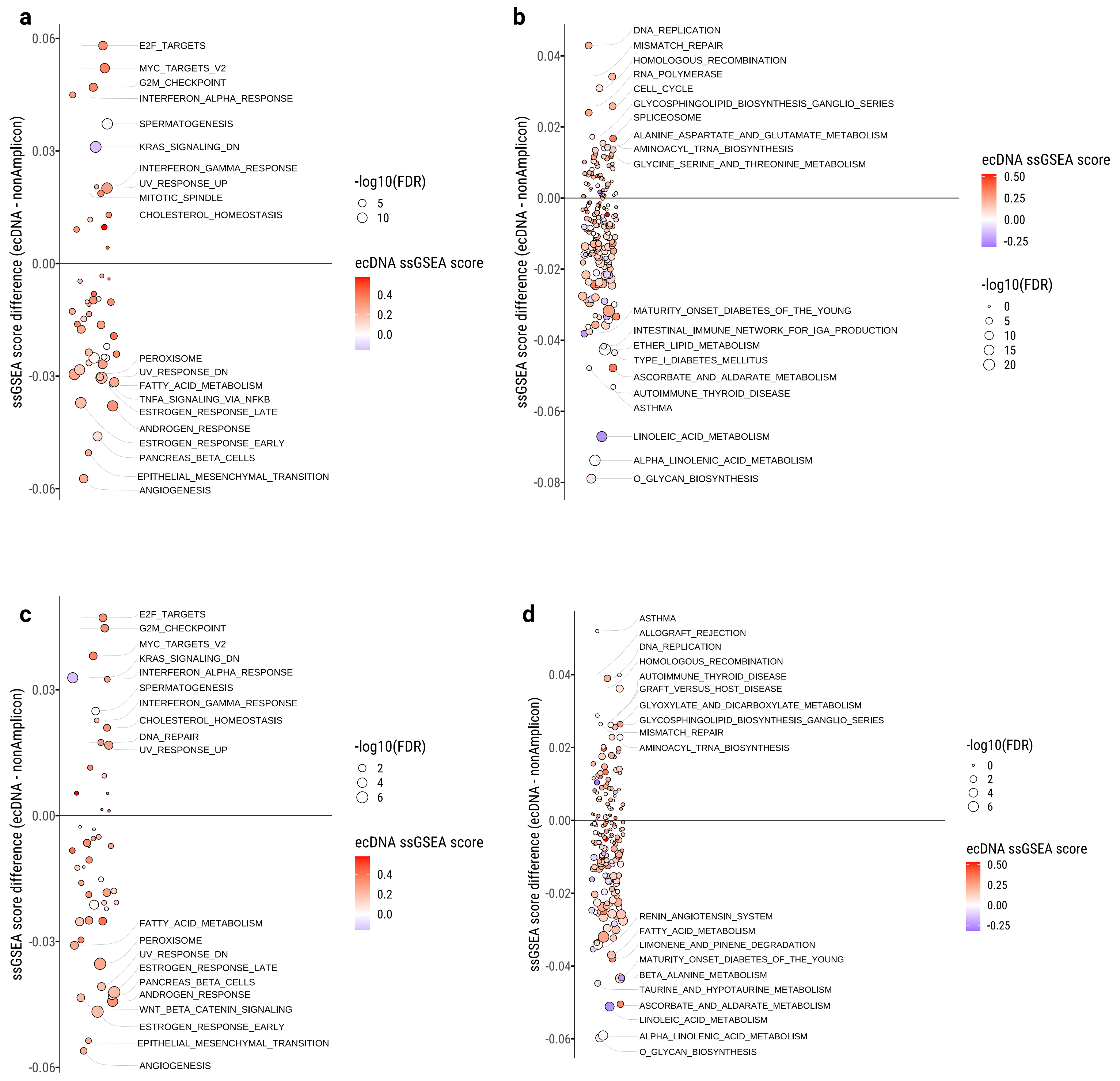

#### Supplementary Fig. 7

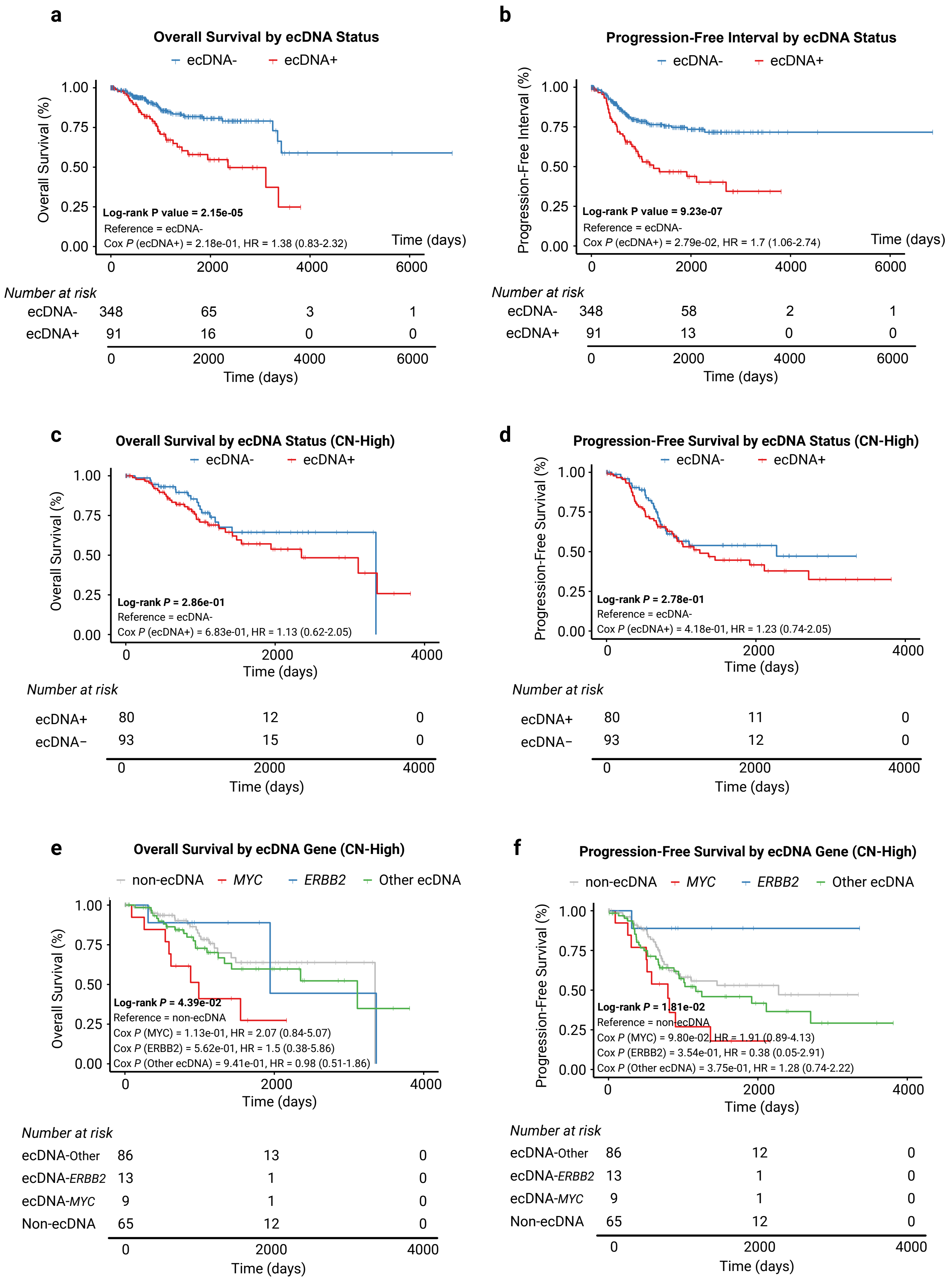

**Supplementary Fig. 8**

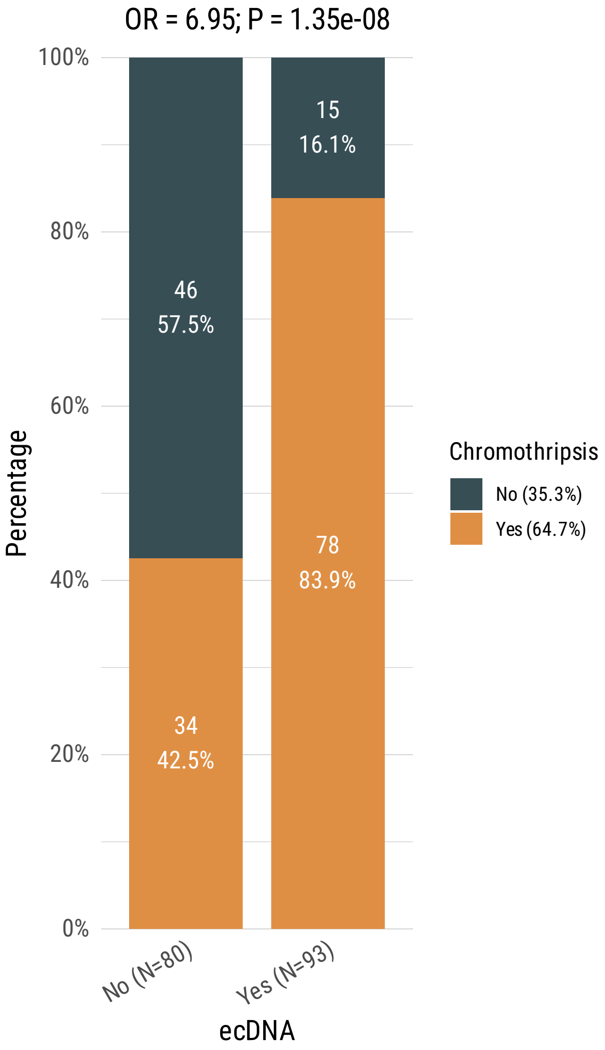

#### Supplementary Fig. 9

a

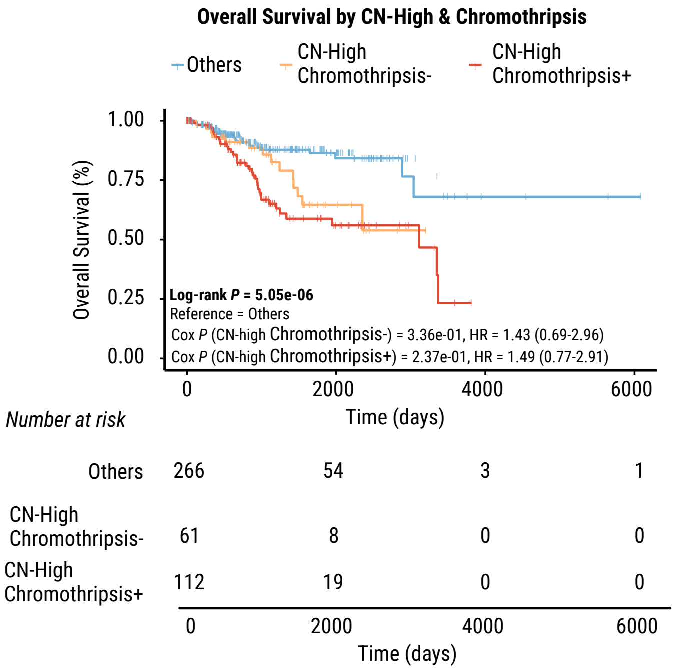


b

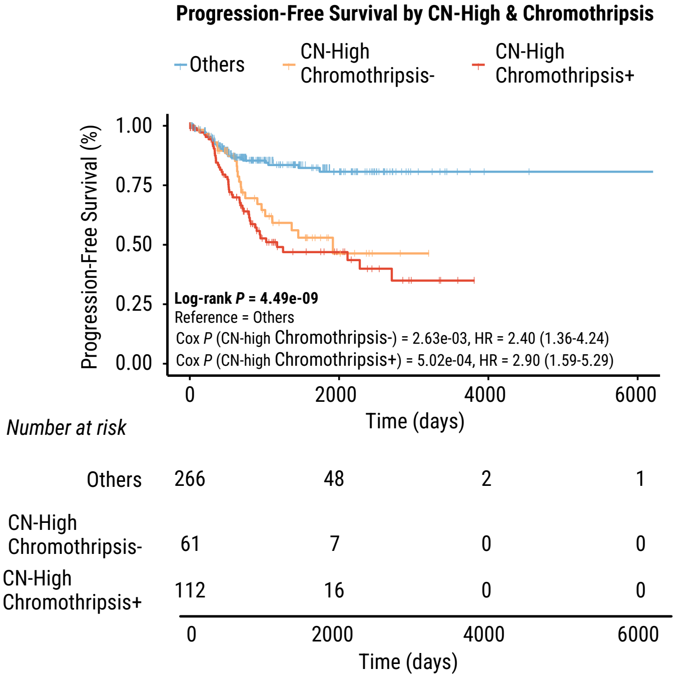

**Supplementary Fig. 10**

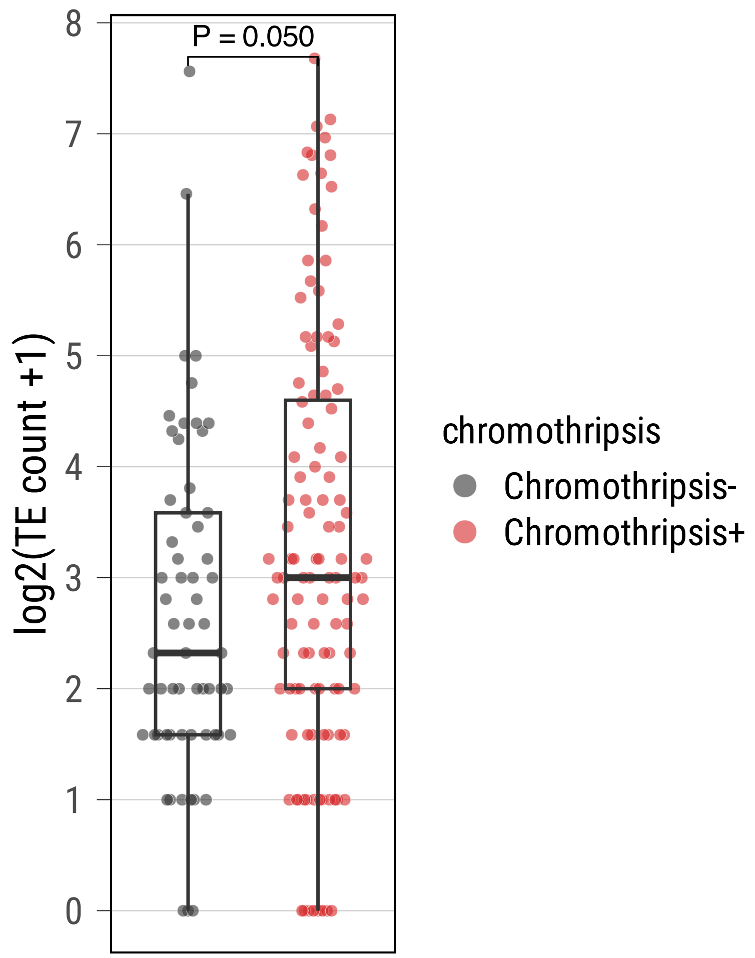

#### Supplementary Fig. 11

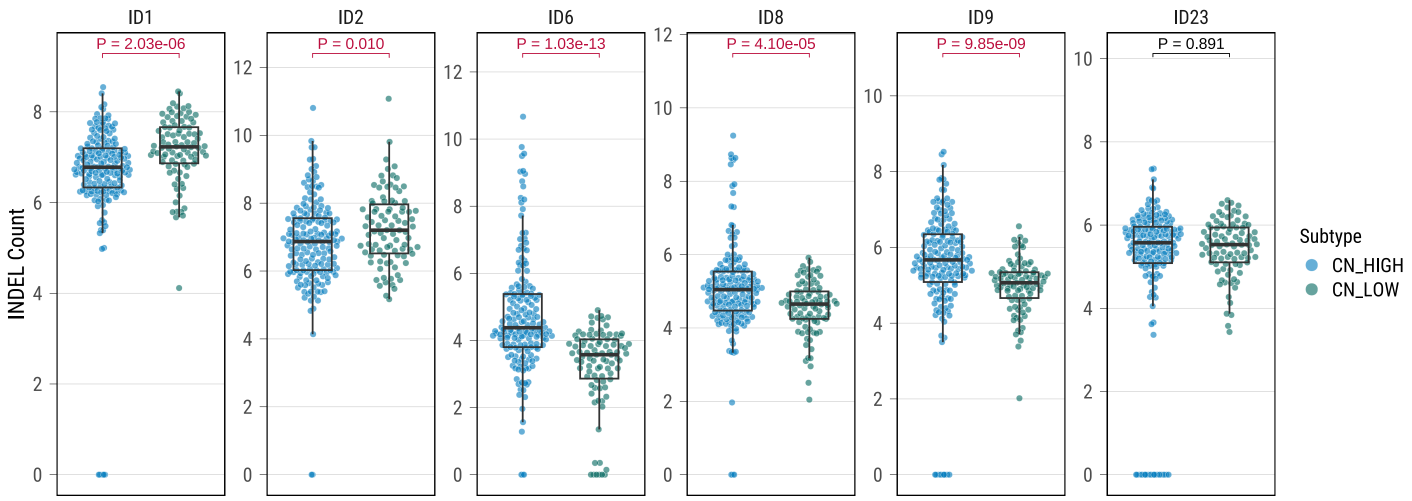

**Supplementary Fig. 12**

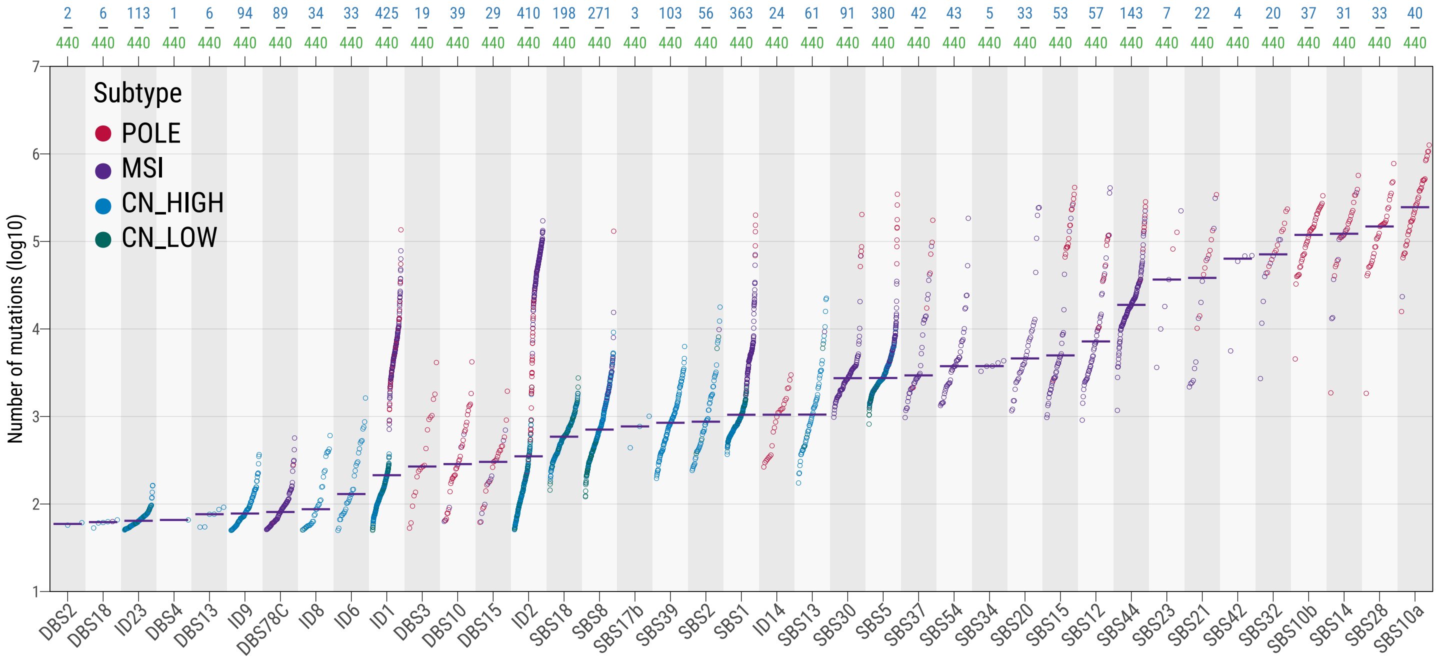

#### Supplementary Fig. 13

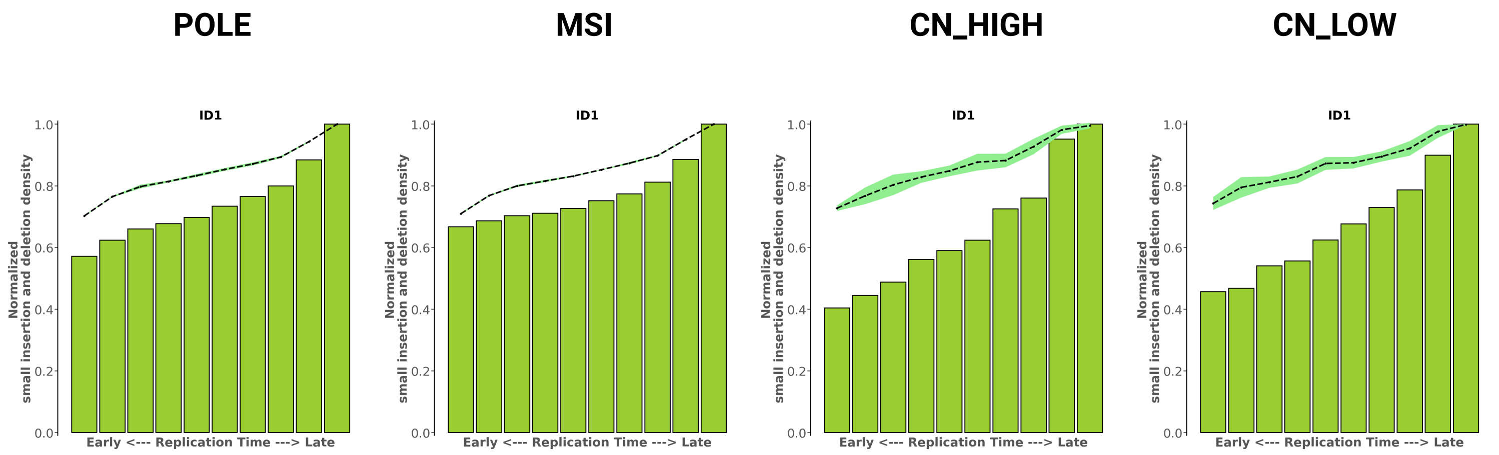

**Supplementary Fig. 14**

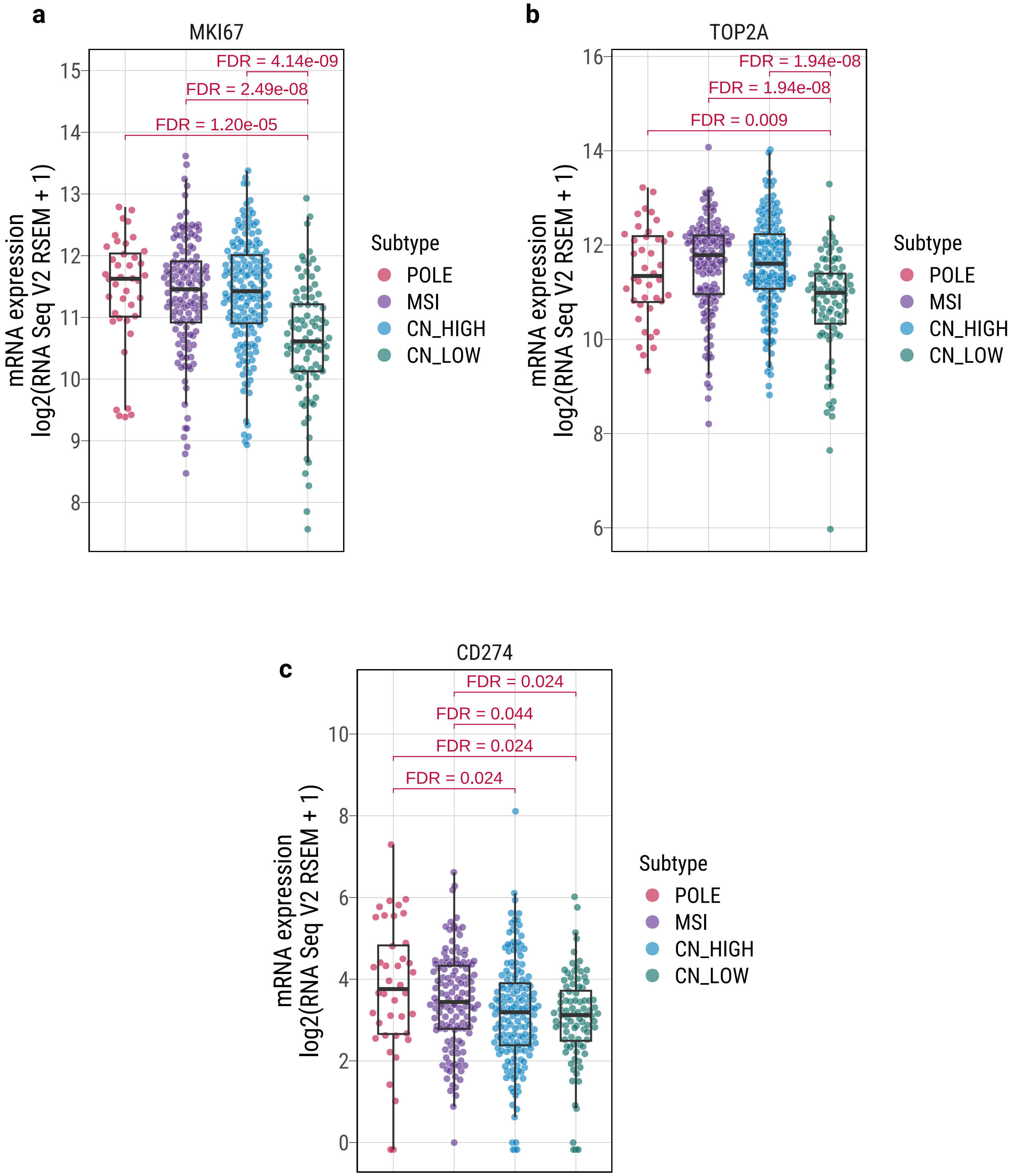

**Supplementary Fig. 15**

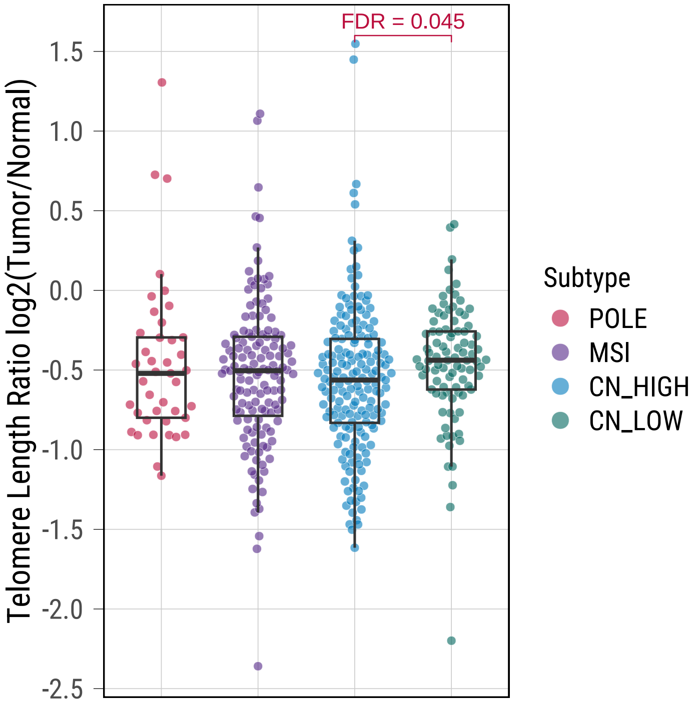

**Supplementary Fig. 16**

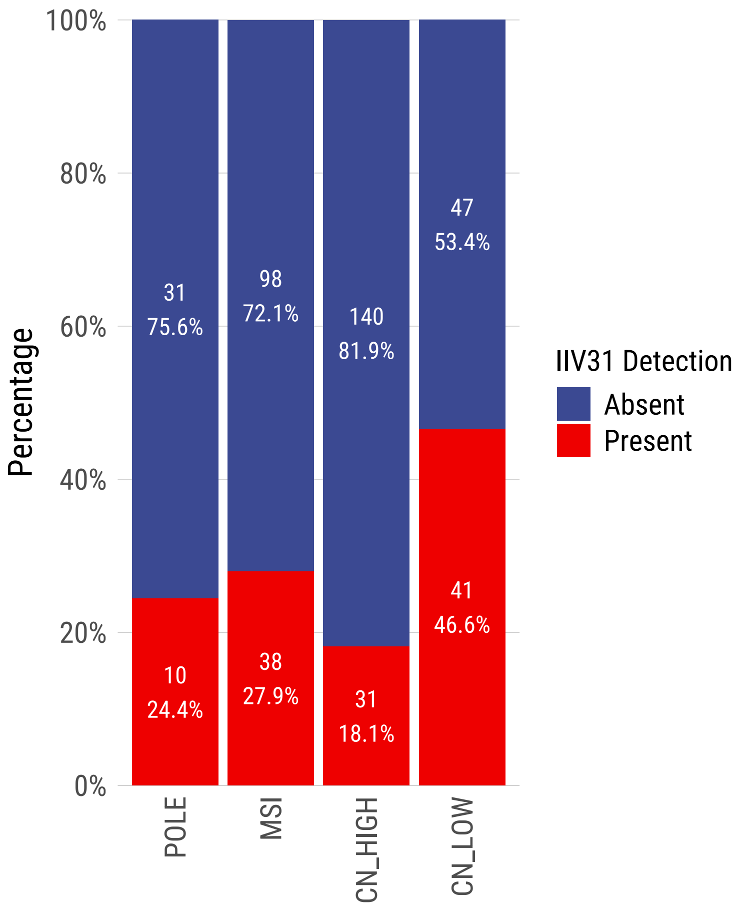

**Supplementary Fig. 17**

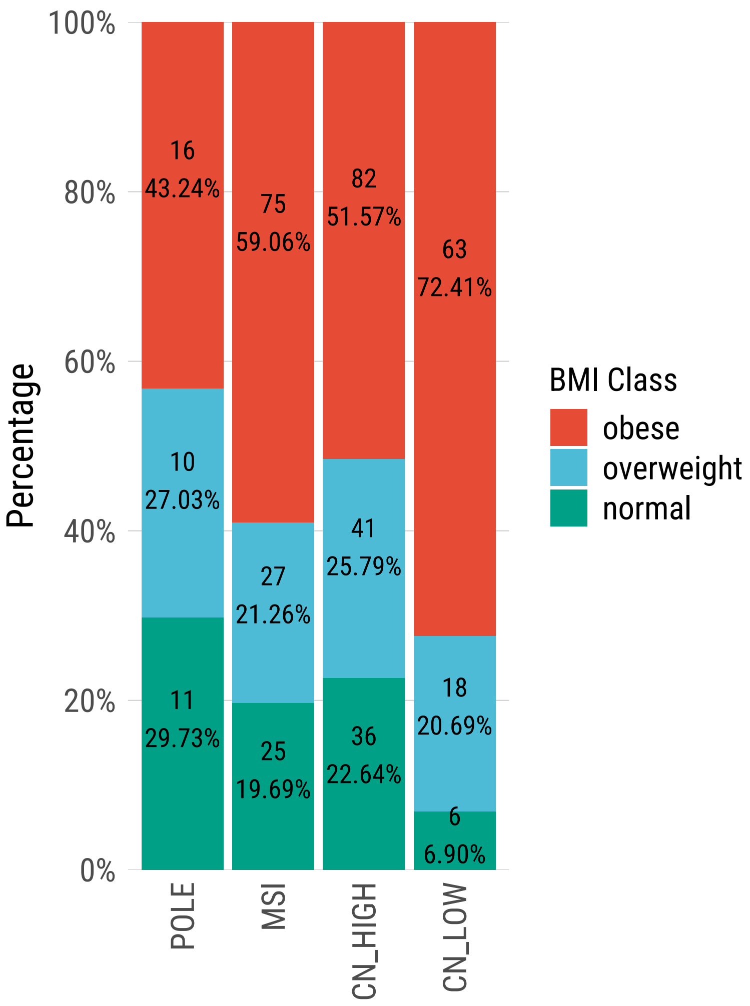

**Supplementary Fig. 18**

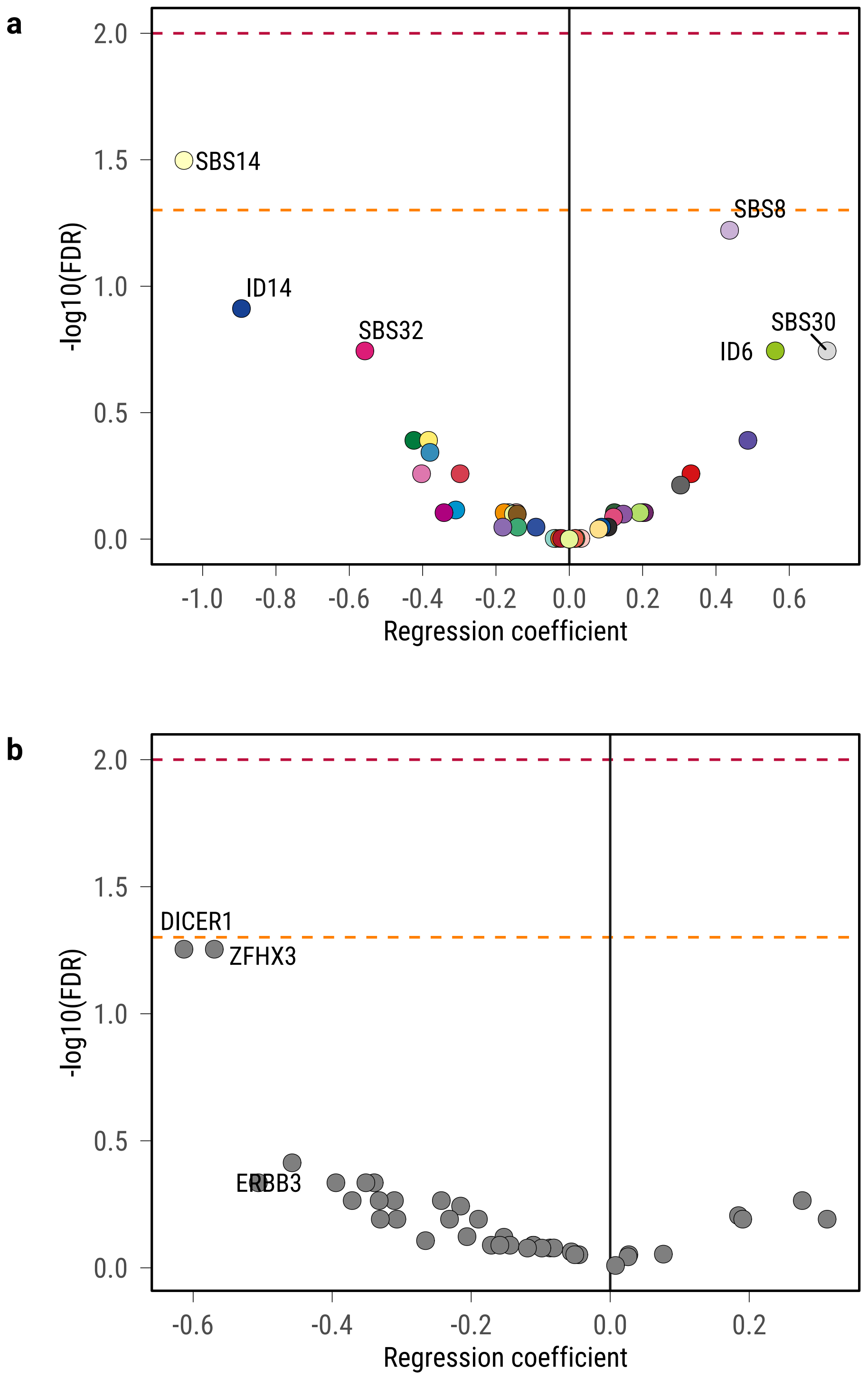
